# FMRP DIRECTLY INTERACTS WITH R-LOOP AND SHOWS COMPLEX INTERPLAY WITH THE DHX9 HELICASE

**DOI:** 10.1101/2021.04.21.440759

**Authors:** Arijita Chakraborty, Arijit Dutta, Leonardo G. Dettori, Jing Li, Leticia Gonzalez, Xiaoyu Xue, Heidi Hehnly, Patrick Sung, Alaji Bah, Wenyi Feng

## Abstract

Mutations in, or deficiency of, FMRP is responsible for the Fragile X syndrome (FXS), the most common cause for inherited intellectual disability. FMRP is a nucleocytoplasmic protein, primarily characterized as a translation repressor with poorly understood nuclear function(s). We recently uncovered a genome protective role of FMRP. We reported that FXS patient-derived cells lacking FMRP sustain higher level of DNA double-strand breaks than normal cells, a phenotype further exacerbated by DNA replication stress. The stress-induced DSBs occur at sequences prone to form R-loops, which are co-transcriptional RNA:DNA hybrids that have been associated with genome instability. Concordantly, we showed that FXS cells accumulate R-loops under replication stress. Moreover, expression of FMRP and not a mutant deficient in binding nucleic acids and known to cause FXS, FMRPI304N, reduced R-loop-associated DSBs. These observations demonstrated that FMRP promotes genome integrity by preventing R-loop accumulation and chromosome breakage. Here, we explore the mechanism through which FMRP prevents R-loop accumulation in an isogenically controlled CRISPR KO of *FMR1* (gene encoding for FMRP) in HEK293T cells. We demonstrate for the first time that FMRP directly binds R-loops. We show that FMRP interacts with DHX9, an RNA helicase that unwinds both double strand RNA and RNA:DNA hybrids and regulates R-loop formation through modulating these activities. This interaction is reduced with FMRPI304N, suggesting that FMRP regulation of R-loop is mediated through DHX9. Interestingly, we show that FMRP inhibits DHX9 helicase activity on RNA:DNA hybrids. Moreover, DHX9 binds chromatin containing R-loops more efficiently in the absence of a functional FMRP. These results suggest an antagonistic relationship between FMRP and DHX9 at the chromatin, where FMRP prevents R-loop formation by suppressing DHX9. Our study sheds new light on our understanding of the genome functions of FMRP.

## INTRODUCTION

Fragile X syndrome (FXS) is a neurodevelopmental disorder due to epigenetic silencing or loss-of-function mutations of the *FMR1* gene encoding FMRP (Ciaccio et al., 2017; Sitzmann et al., 2018). FMRP is a nuclear-cytoplasmic RNA binding protein that regulates multiple biological processes of its diverse mRNA substrates, including their maturation in the nucleus, nuclear export, cytoplasmic transport, and ultimately, their translation at the synapse (Banerjee et al., 2018; Sudhakaran et al., 2014; Zhou et al., 2017). The ability of FMRP to participate in multiple processess in the cell is attributed to the presence of multiple domains and their relative 3D-organization. All FMRP splice variants contain two amino (N-)terminal methylated lysine-binding Agenet domains (Age1 and Age2), three K-homology (KH0, KH1, and KH2) RNA binding domains and a highly variable (isoform-specific) carboxy (C-)terminal intrinsically <u>d</u>isordered <u>r</u>egion (C-IDR), which in the case of the predominant isoform 1, contains an RNA binding RGG-box. Additionally, the presence of a nuclear localization signal and a nuclear export signal allows FMRP to shuttle between the nucleus and the cytoplasm, with approximately 4% of FMRP detected in the nucleus (Feng et al., 1997b).

Though its exact role in the nucleus is not clear, recent studies have suggested that FMRP is also involved in genome maintenance (Dockendorff and Labrador, 2019). We recently demonstrated that FXS patient-derived cells accumulate genome-wide DNA double strand breaks (DSBs), particularly during replication stress (Chakraborty et al., 2020). We further demonstrated that the DSBs in FXS cells were associated with R-loops (Chakraborty et al., 2020), which are three-stranded nucleic acid structures formed during transcription when the nascent RNA stably anneals to the template DNA strand, displacing the non-template DNA strand (Thomas et al., 1976). R-loops play important roles in gene expression and many biological processes, but they are also an important source of genomic instability, particularly when R-loop formation is exacerbated by replication-transcription conflict (Crossley et al., 2019; Garcia-Muse and Aguilera, 2019). Consequently, there are an abundance of cellular proteins that interact with R-loops and promote their resolution. These include helicases that unwind the RNA:DNA hybrids within the R-loop structure, topoisomerases that release the negative supercoil in the DNA duplex behind the transcription machinery, and ribonucleases that degrade the RNA from the RNA:DNA hybrids. In addition, many other regulatory factors have been associated with R-loop metabolism.

We show that expression of FMRP, but not the FMRP-I304N mutant, ameliorated DSB formation induced by such conflict (Chakraborty et al., 2020). Thus, our work suggested a genome protective role of FMRP by preventing R-loop accumulation during replication-transcription conflict. However, several key questions remain. Does FMRP interact with R-loop and if so, how does it promote R-loop resolution? Does it interact with the R-loop resolution factors mentioned above? Here we investigated how FMRP promotes R-loop resolution by testing if FMRP directly interacts with R-loops and R-loop resolvases. We purified recombinant FMRP and measured its capacity to bind various nucleic acid structures by electrophoretic mobility shift (EMSA) assay. We present evidence that FMRP interacts directly with R-loop structures specifically through its C-IDR, making FMRP the archetype of a class of IDR-based R-loop “reader” proteins (Dettori et al., 2021). We also present evidence of FMRP co-immunoprecipitating with known R-loop regulator proteins including Top3β and DHX9, suggesting that FMRP might mediate the interaction between these proteins and R-loop structures. In this study, we focused on the characterization of the interplay between FMRP and DHX9, an RNA helicase known to unwind R-loops. DHX9 has been reported to have apparently opposing functions during R-loop regulation. On one hand, DHX9 knockdown HEK293T cells showed increased R-loop formation, suggesting that DHX9 prevents R-loop accumulation (Cristini et al., 2018). Consistent with this observation it was recently shown that DHX9 is recruited by the TDRD3/ Top3β complex to remove R-loops at specific target genes (Yuan et al., 2021). On the other hand, in cells depleted of the SFPQ RNA splicing protein the loss of DHX9 led to reduced R-loop levels, suggesting that DHX9 in fact promotes R-loop formation by unwinding dsRNA when RNA splicing is impaired (Chakraborty et al., 2018). Thus, these studies suggest that DHX9 might prevent or promote R-loop formation in different genetic contexts, making it a challenging but also important target to study the complex nature of R-loop regulation.

Here we present evidence that FMRP directly interacts with DHX9 and regulates its helicase activity, subcellular localization, and ultimately chromatin association. Contrary to our original hypothesis that FMRP recruits DHX9 to R-loop within specific gene substrates to facilitate R-loop resolution, we observed that FMRP inhibits DHX9 helicase activity *in vitro* and chromatin R-loop association *in vivo.* These unexpected results led us to propose a model that in the chromatin context FMRP serves as a signal, through protein-protein interaction, for DHX9 to disengage from the R-loop after it unwinds the RNA:DNA hybrid. Our study represents a significant advance in the understanding of the mechanisms by which FMRP regulates an R-loop resolution enzyme and promotes genome integrity upon replication stress.

## RESULTS

### FMRP is enriched in the nucleus and co-localizes with R-loops in response to DNA replication stress in human lymphoblastoid cells

We previously showed that FXS patient-derived cells lacking FMRP have elevated genome-wide DSBs near R-loop forming sites when undergoing replication stress by aphidicolin (APH), a DNA polymerase inhibitor (Chakraborty et al., 2020). We proposed that FMRP protects the genome by preventing DSBs during induced replication-transcription conflict. Here we asked whether FMRP alters its expression level and/or its cellular localization in response to APH (Figure 1A). First, the total level of FMRP remained the same with and without APH (Figure S1A). However, the nuclear fraction of FMRP increased from 18% in DMSO (vehicle)-treated control cells to 24-36% in APH treatment (Figure S1B). In contrast, GAPDH (cytoplasmic) and Histone H3 (nuclear) controls maintained their respective subcellular localization, with or without APH (Figure S1B). We concluded that FMRP has substantial nuclear fraction in human lymphoblastoids, and it becomes further enriched in the nucleus in response to replication stress. Next, we wanted to visualize the localization of FMRP relative to R-loops. Immunofluorescence microscopy in lymphoblastoid cells revealed a distinct staining pattern of FMRP, which was predominantly distributed in the cytoplasm and at the periphery of the nucleus (Figure 1B & Figure S1C) in untreated and DMSO-treated cells. Upon induction with APH, FMRP was enriched in the nucleus, consistent with the chromatin fractionation experiments. RNA:DNA hybrid signals as observed from S9.6 antibody staining were present in both cytoplasm and the nucleus, and significantly induced under APH treatment (Figure 1B, C & Figure S1C, D). To determine the specificity of the staining we treated the cells with RNase H, which degrades RNA:DNA hybrids. Indeed, it significantly reduced the RNA:DNA hybrid signals specifically in APH treated samples (Figure 1C). In addition, we treated the cells with RNase III in order to eliminate the possibility that S9.6 was non-specifically targetting dsRNA (Smolka et al., 2021). Even though, S9.6 was reduced significantly upon RNase III treatment, a pattern of increased R-loops was maintained in APH and was significantly higher than the untreated/DMSO samples (Figure S1D). This pattern was lost in the RNase H treatment (Figure 1C). Altogether, these results indicate enhanced R-loop formation with APH which is in line with our previous observation that APH induces DNA DSBs even in normal cells (Chakraborty et al., 2020). Moreover, FMRP signals were closely associated with the RNA:DNA hybrid signals. Quantification of co-localization of the two signals indicated that the percentage of FMRP overlapping with RNA:DNA hybrid signals increases upon drug treatment (Figure 1D & Figure S1E). Notably, this co-localization is reduced in RNase H treatment of the cells in comparison to its mock but remains unchanged upon RNase III treatment relative to only buffer, suggesting that the co-localization thus observed is specific for FMRP and RNA:DNA hybrids.

**Figure 1.**
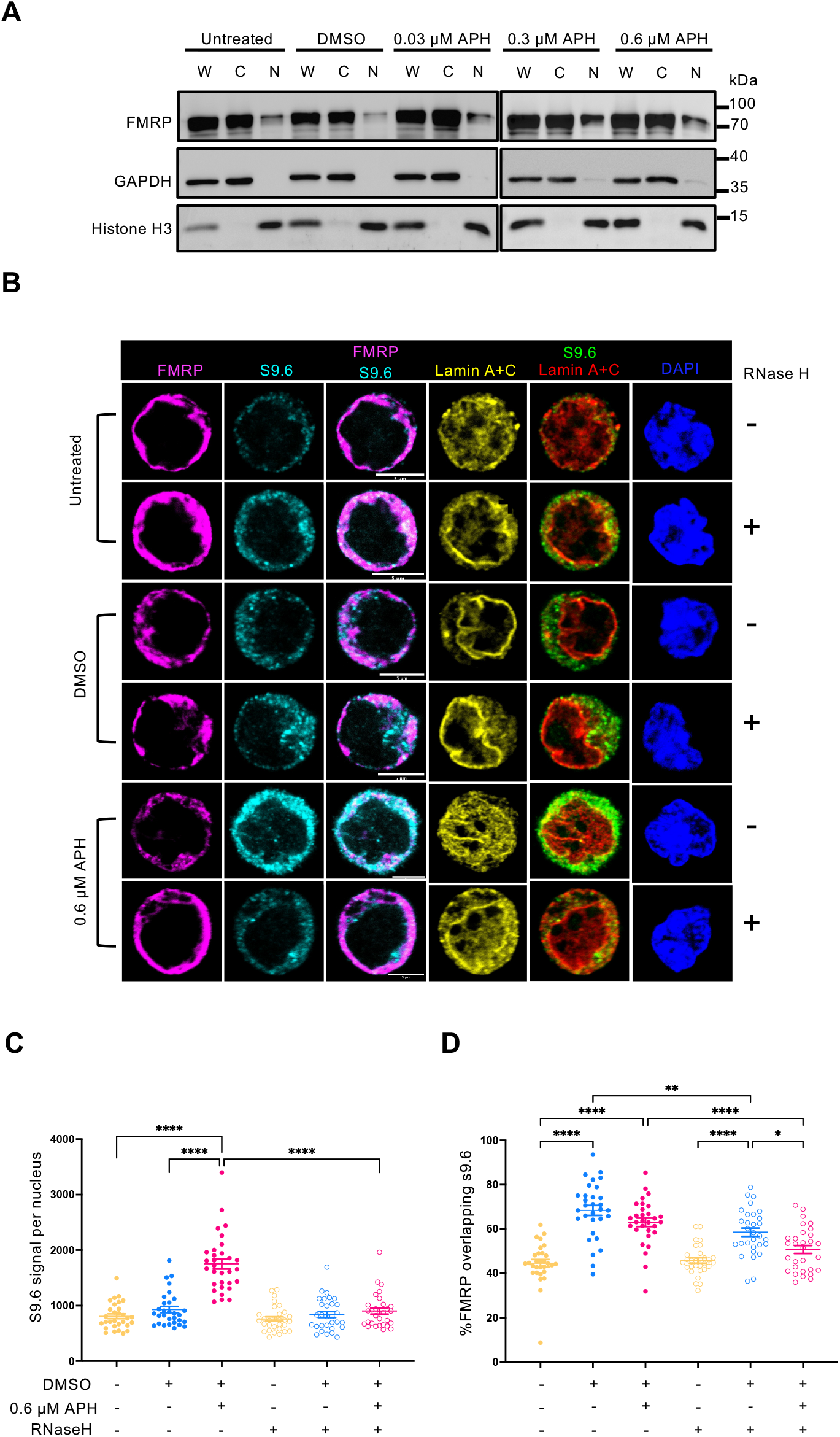
FMRP is enriched in the nucleus upon replication stress and co-localizes with R-loops. **(A)** Subcellular fractionation of FMRP. Western blot showing, whole cell extract (W), cytoplasmic fraction (C) and nuclear fraction (N) of lymphoblastoid cells from unaffected control (NM) with and without replication stress. GAPDH and Histone H3 serve as cytoplasmic and nuclear controls, respectively. Two independent experiments were conducted, and one representative experiment is shown. **(B)** Co-localization of FMRP and RNA:DNA hybrids. Immunofluorescence images of untreated, DMSO and APH treated GM6990 cells co-stained for RNA:DNA hybrids (cyan), FMRP (magenta), lamin A/C (yellow) and DAPI (blue). Cells were treated with RNase H enzyme to show specificity for staining R-loops by S9.6 antibody. Immuno-staining is shown in a single Z-plane. Scale bar, 5 µm**. (C)** Quantification of S9.6 signal per nucleus in cells treated with or without RNase H. Error bars indicate standard error of mean (SEM),N∼30 cells. One-way ANOVA followed by Tukey’s multiple comparison test, *p < 0.05, **p<0.01, ****p < 0.0001. **(D)** Quantification of colocalization of FMRP with S9.6 signal using Coloc2 plugin. Error bars indicate SEM,N∼30 cells per sample. One-way ANOVA followed by Tukey’s multiple comparison test, *p < 0.05, **p<0.01, ****p < 0.0001.

### FMRP directly binds R-loops through its C-IDR and the interaction is weakened by the I304N mutation in the KH2 domain

The observed colocalization described above suggests a potential interaction between the protein and R-loops. Therefore, to test the ability of FMRP to directly bind R-loops, we resorted to recombinantly expressing and purifying full length FMRP, the N-terminal folded domain (N-Fold) and the C-IDR (Figure 2A and Figure S2A&C). We then measured their binding affinities for R-loops with and without RNA overhang and R-loop sub-structures including ssDNA, dsDNA, RNA, and DNA:RNA hybrid (Figure 2B) in an electrophoretic mobility shift assay (EMSA). DNA:RNA hybrids with or without a 5’ DNA overhang produced nearly identical results for all proteins and therefore only DNA:RNA without overhang is shown. First, we observed binding between both the N-Fold and C-IDR of FMRP to the R-loop with 5’ RNA overhang and the aformentioned sub-structures of R-loops with varying affinities (Figure 2C&D, Figure S3 and Table 1). Due to the high propensity of FMRP to aggregate and precipitate at high concentrations (Sjekloca et al., 2009; Sjekloca et al., 2011), it was not feasible to obtain complete binding isotherms and determine the dissociation constants (K_D_) for some weak FMRP:substrate interactions (Table 1). Of all the tested protein-nucleic acid pairs, the C-IDR and R-loop without overhang showed the highest affinity (K_D_ = 4.73±3.8 nM, Figure 2E, Figure S3 and Table 1). Intriguingly, the interaction was weakened with a 5’ RNA overhang to the R-loop (K_D_ = 148.3±10.3 nM, Figure 2D). Moreover, while the C-IDR showed affinity towards ssDNA and dsDNA in isolation, it barely interacted with the DNA:RNA hybrid or ssRNA (Figure 2H Figure S3 and Table 1). We note that the lack of binding for ssRNA might be due to the substrate lacking any consensus FMRP binding motifs. Therefore, we concluded that the C-IDR interacted with R-loops through simultaneous binding to the ssDNA and dsDNA junction, with the RNA overhang interfering with the interaction. In contrast, the N-Fold bound R-loops with ssRNA overhang more tightly than those without overhang, albeit with still lower affinity than C-IDR (Figure 2F, Figure S3 and Table 1). Additionally, N-Fold showed affinities for ssRNA and ssDNA, but not dsDNA nor the DNA:RNA hybrid (Figure 2F, Figure S3 and Table 1). Therefore, the N-Fold likely interacts with the R-loop through binding with the single stranded segments (RNA or DNA) of the R-loop. Finally, the affinity of the full length FMRP for R-loop with or without overhang (K_D_ = 615.1±7.1 nM and 2398±2.8 nM, respectively) was markedly decreased compared to the C-IDR (Figure 2I&J, Figure S3 and Table 1). We surmised that in the full length protein the N-Fold actually interferes with the C-IDR, possibly through a long-range intramolecular mechanism, for its binding to R-loop, despite their preference for different substructures of the R-loop. Thus, our results demonstrated that the FMRP binding to R-loops involves multivalent interactions, with N-Fold and C-IDR showing varying affinities to all segments of an R-loop structure. Moreover, these multivalent interactions between FMRP and R-loops are modulated by intra-and inter-molecular cooperative and/or inhibitory effects within FMRP, as well as between FMRP and the R-loop sub-structures.

**Figure 2.**
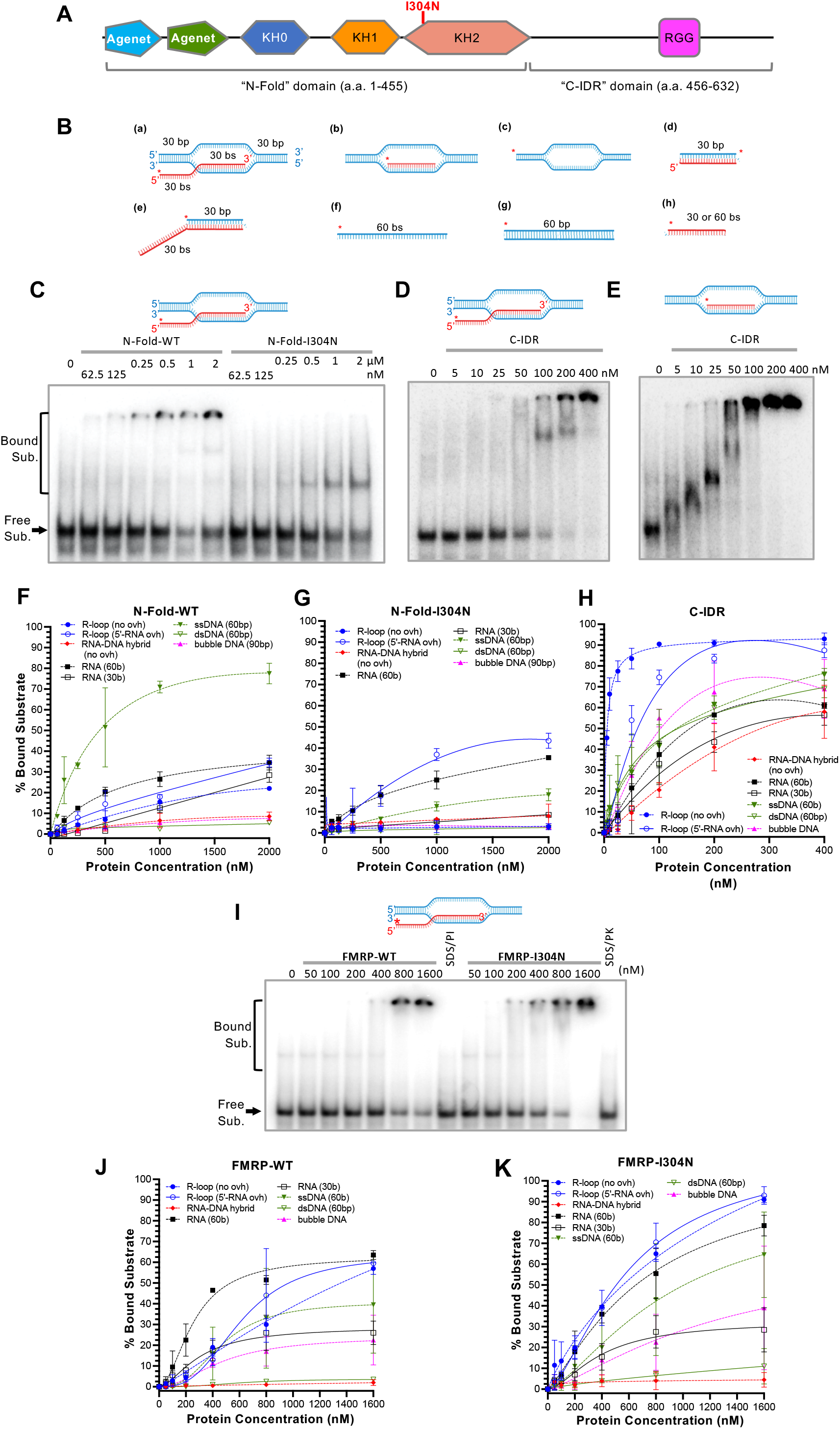
FMRP directly binds R-loops *in vitro.* **(A)** Schematic representation of FMRP protein domains, indicating the fold region and the C-terminus intrinsically disordered region (C-IDR). The folded FMRP domain also harbor the isoleucine to asparagine mutation at residue 304 which causes FXS. **(B)** Nucleic acid structures used in the electrophoretic mobility shift assay (EMSA) to determine binding interaction with FMRP N-Fold domain or FMRP C-IDR. Blue strand represents DNA and red represents RNA, while asterisk indicates P^32^ label at the 5’-end of the DNA or RNA strand. a- R-loop with 5’-RNA overhang (5’-RNA ovh), b- R-loop with no overhang (no ovh), c- Bubble DNA (90 bp), d- RNA:DNA hybrid (no ovh), e- RNA:DNA hybrid (5’-RNA ovh), f- Single-stranded DNA (ssDNA), g- Double-stranded DNA (dsDNA) and h- RNA (30 or 60 bs). **(C&D)** Representative EMSAs for interaction between R-loop with 5’-RNA overhang and the N-Fold and C-IDR domains. Sub., substrates. **(E)** Representative EMSA for interaction between R-loop without overhang and the C- IDR. **(F-H)** Quantification of the percentage of bound nucleic acid substrates at the indicated protein concentrations for N-Fold-WT **(F),** N-Fold-I304N **(G)** and C-IDR **(H). (I)** Representative EMSA for interaction between R-loop with 5’-RNA overhang and the full length FMRP with or without I304N mutation. **(J&K)** Quantification of the percentage of bound nucleic acid substrates at the indicated protein concentrations for FMRP-WT (J) and FMRP-I304N (K). The free and bound substrates labeled for **(C)** is true for all EMSA gels.

**Table 1.** Dissociation constants (K_D_) of FMRP domains for nucleic acid substrates.

| **Substrate** | **FMRP-WT** | **FMRP-I304N\*** | **N-Fold-WT** | **N-Fold-I304N** | **C-IDR** |
| --- | --- | --- | --- | --- | --- |
| R-loop (no ovh) | 2398.0 ± 2.8 | 1753.0 ± 5.5 | NA | NA | 4.7 ± 3.8 |
| R-loop (5' ovh) | 615.1 ± 7.1 | 594.7 ± 7.1 | 320 ± 3.0 | NA | 148.3 ± 10.0 |
| DNA bubble | 471.6 ± 4.4 | 1276.0 ± 10.2 | NA | NA | 346.4 ± 8.7 |
| dsDNA (60 bp) | 602.9 ± 0.3 | NA | 41.98 ± 1.0 | NA | 56.8 ± 7.1 |
| RNA (30 bs) | 472.2 ± 10.5 | 920.7 ± 11.9 | NA | NA | 837.1 ± 5.9 |
| RNA (60 bs) | 260.3 ± 4.4 | 720.0 ± 5.2 | 615.9 ± 2.2 | NA | 649.9 ± 3.2 |
| ssDNA (60 bs) | 302.4 ± 3.9 | 388.5 ± 4.9 | 616.9 ± 7.9 | NA | 79.59 ± 9.3 |
| RNA-DNA hybrid | NA | NA | NA | NA | NA |

$K_D$ values were calculated from averaging two independent measurements for every protein/substrate combination. $K_D$ values in blue are those with calculable confidence intervals, in contrast to those in black for which a confidence interval could not be calculated. "NA", $K_D$ values could not be calculated. "ovh", overhang.

Next, we investigated the effect on R-loop binding by I304N, an FXS-causing mutant defective in RNA binding and polysome association (De Boulle et al., 1993; Feng et al., 1997a). We recently showed that FMRP-I304N had reduced ability to suppress R-loop-induced DSBs during programmed replication-transcription conflict (Chakraborty et al., 2020). We generated both the full length FMRP and the N-Fold containing the I304N substitution (Figure 2A and Figure S2B&D). The mutation indeed disrupted the interactions between the N-Fold and all substrates tested (Figure 2C&G, Figure S3 and Table 1). In contrast, the I304N mutation caused the full length FMRP to bind R-loops with or without overhang at moderately higher affinity, with a K_D_ of 594.7±7.1 nM and 1753±5.5 nM, respectively (Figure 2I, Figure S3 and Table 1). However, since FMRP lacks apparent protein domains for helicase or nuclease activitiy, we surmised that its ability to resolve R-loops must come from its association with its binding proteins. Therefore, we next tested if FMRP interacts with known R-loop-interacting proteins. At the beginning of our study few FMRP-binding proteins with functions in the R-loop pathway existed in the literature, so we exploited a large-scale human proteome study and collected all reported interactions with FMRP (Hein et al., 2015). Both FMRP and DHX9 were pulled down by THOC1, a component of the THO nuclear export complex. Depletion of the THO complex causes DNA damage that is R-loop dependent (Dominguez-Sanchez et al., 2011). DHX9 is an RNA helicase known to suppress R-loop formation and prevent DNA DSBs (Chakraborty and Grosse, 2011; Cristini et al., 2018). Therefore, we set out to investigate the potential interaction between FMRP and DHX9.

### FMRP interacts with DHX9 and the interaction is at least partially dependent on a functional KH2 domain and R-loop formation

Using the aforementioned GM06990 lymphoblastoids we first demonstrated co-immunoprecipitation (IP) of FMRP and its known interacting protein, FXR1 (FMR1 autosomal homolog 1), as a positive control (Zang et al., 2009) (Figure S4D). We also detected DHX9 interaction with FMRP through co-IP (Figure S4E). In addition, the complex pulled down by anti-DHX9 also comprised of Top3β (Figure S4E), which has been implicated in R-loop suppression by reducing negatively supercoiled DNA behind RNA polymerase II (Yang et al., 2014).

We then asked if the *in vivo* FMRP-DHX9 interaction is (i) mediated by non-specific RNA binding, (ii) dependent on R-loop formation and (ii) dependent on the KH2 domain of FMRP. To address these questions we first generated a CRISPR knock-out (KO) of *FMR1* in HEK293T cells (Figure 3A). We selected a *fmr1*KO-B3 clone and showed that it modeled the elevated DNA damage feature of the FXS patient-derived cells we previously described (Figure 3B-D) (Chakraborty et al., 2020). We then generated stable cell lines expressing enhanced green fluorescent protein (eGFP)-tagged FMRP and FMRP-I304N in the *fmr1*KO cells to facilitate the comparison of wild type to mutant FMRP with respect to their interaction with DHX9. Our analyses demonstrated that the *fmr1KO* cells expressing eGFP-FMRP significantly reduced APH-induced DSBs and R-loop formation compared to cells expressing only eGFP, or the mutant eGFP-FMRP-I304N (Figure 3E&F).

**Figure 3.**
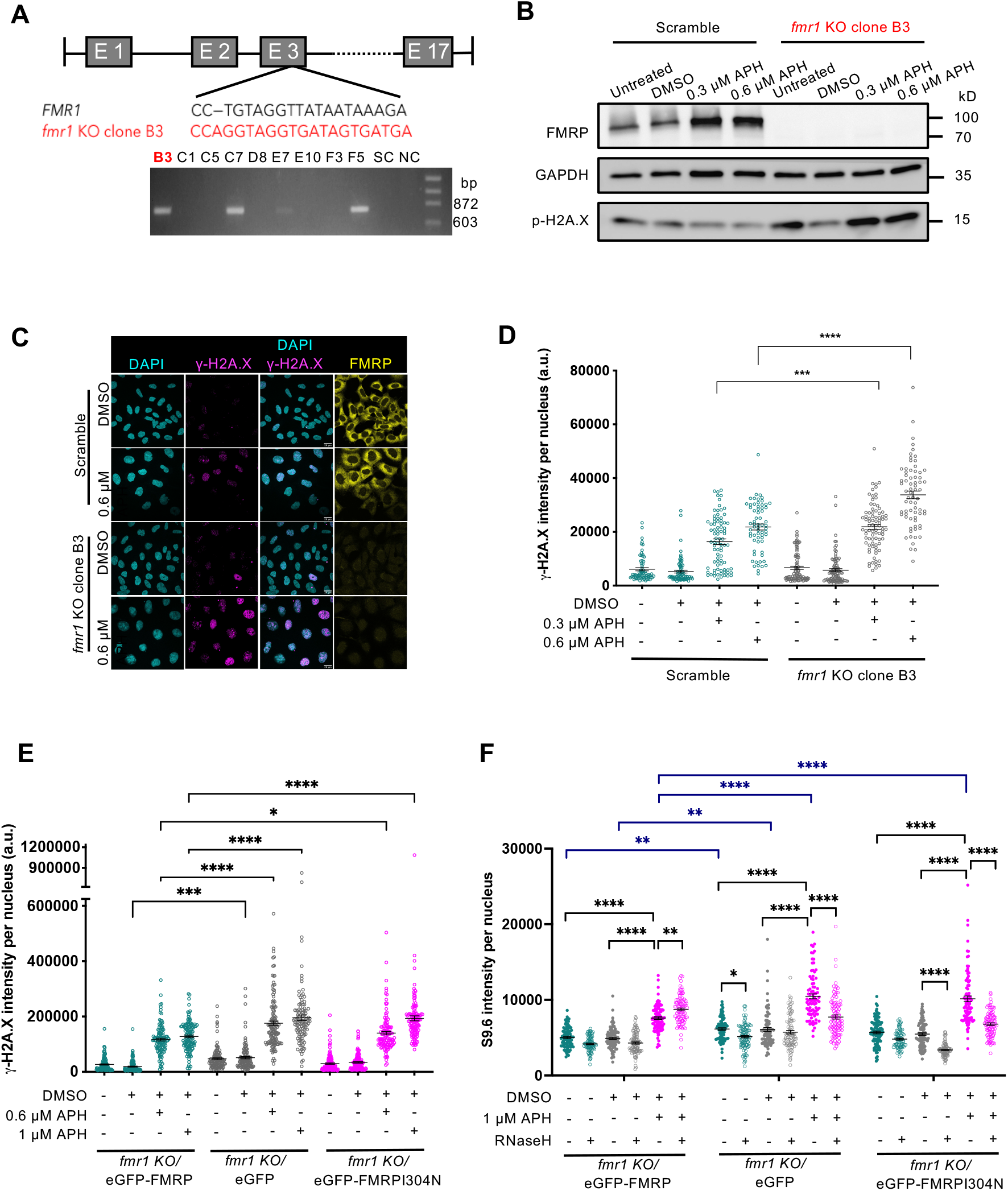
CRISPR KO of *FMR1* gene in HEK293T cells and re-expression of FMRP. **(A)** Genome structure of *FMR1* with CRISPR target region in exon 3. CRISPR clones analyzed by PCR. Clone B3 is used for all subsequent experiments. “SC”, scramble; “NC”, no template control. **(B)** Western blot confirms the lack of FMRP expression in *fmr1* KO cells. **(C&D)** Increased DNA damage by APH-induced replication stress in *fmr1* KO cells. Scale bar, 20 µm. N>60 cells per sample were analyzed. One-way ANOVA test followed by Tukey’s multiple testing for all pair-wise comparisons was performed. **(E)** Retroviral transduction and stable cell line generation of eGFP, eGFP-FMRP and eGFP-FMRP-I304N expression in *fmr1* KO cells. DNA damage by APH-induced replication stress after re-expressing FMRP, FMRP-I304N or nothing. N>110 cells per sample were analyzed. One-way ANOVA test followed by Sidak’s multiple testing. Two independent experiments were done and a representative experiment is shown for (D) and (E)**. (F)** Elevated S9.6 in eGFP and eGFP-FMRP-I304N expressing *fmr1* KO cells. S9.6 signals were sensitive to RNase H treatment. N> 75 cells were analyzed. Two-way ANOVA test followed by Holm-Sidak’s multiple testing. Annotation for P values are: *, p<0.05; **, p<0.01; ***, p<0.001; ****, p<0.0001. Error bars indicate standard error of mean (SEM).

We carried out reciprocal IP reactions and calculated the ratios of co-immunoprecipitated protein to the immunoprecipitated protein (Co-IP/IP) in each condition. First, we showed that the FMRP-DHX9 interaction is not mediated by non-specific association with RNA (Figure 4A). Second, we indeed observed co-immunoprecipitation in all conditions in cells carrying wild type FMRP, thus confirming *in vivo* interaction between FMRP and DHX9 (Figure 4B, top and bottom panels). Third, the Co-IP/IP ratios decreased in cells expressing RNase HI compared to those expressing a catalytically dead mutant RNase HI (Figure 4B, compare RNase HI “+” to “-”), indicating that the FMRP-DHX9 interaction is at least partially dependent on R-loop. Note that we observed higher ratios of FMRP/DHX9 than DHX9/FMRP in all conditions in cells carrying wild type FMRP. We think this is due to the fact that the α-DHX9 antibody is far more efficient than α-FMRP, making co-IP of FMRP by α-DHX9 more readily detectable. This is reflected in the greater variability of co-IP of DHX9 by α-FMRP. Additionally, FMRP is predominantly cytoplasmic whereas DHX9 is almost exclusively nuclear, suggesting that their interaction most likely occurs in the nucleus, which is also more readily detectable by α-DHX9. Fourth, IP by α-DHX9 pulled down less FMRP-I304N compared to FMRP (compare the FMRP/DHX9 ratios for cells carrying FMRP or FMRP-I304N, Figure 4B, top panel). Similarly, α-FMRP pulled down less DHX9 in cells carrying FMRP-I304N than those carrying FMRP, though the reduction was less significant than the reciprocal IP above (Figure 4B, bottom panel). These results led us to conclude that the *in vivo* interaction between FMRP and DHX9 is at least partially dependent on R-loops and the KH2 domain of FMRP. We next tested if FMRP directly interacts with DHX9.

**Figure 4.**
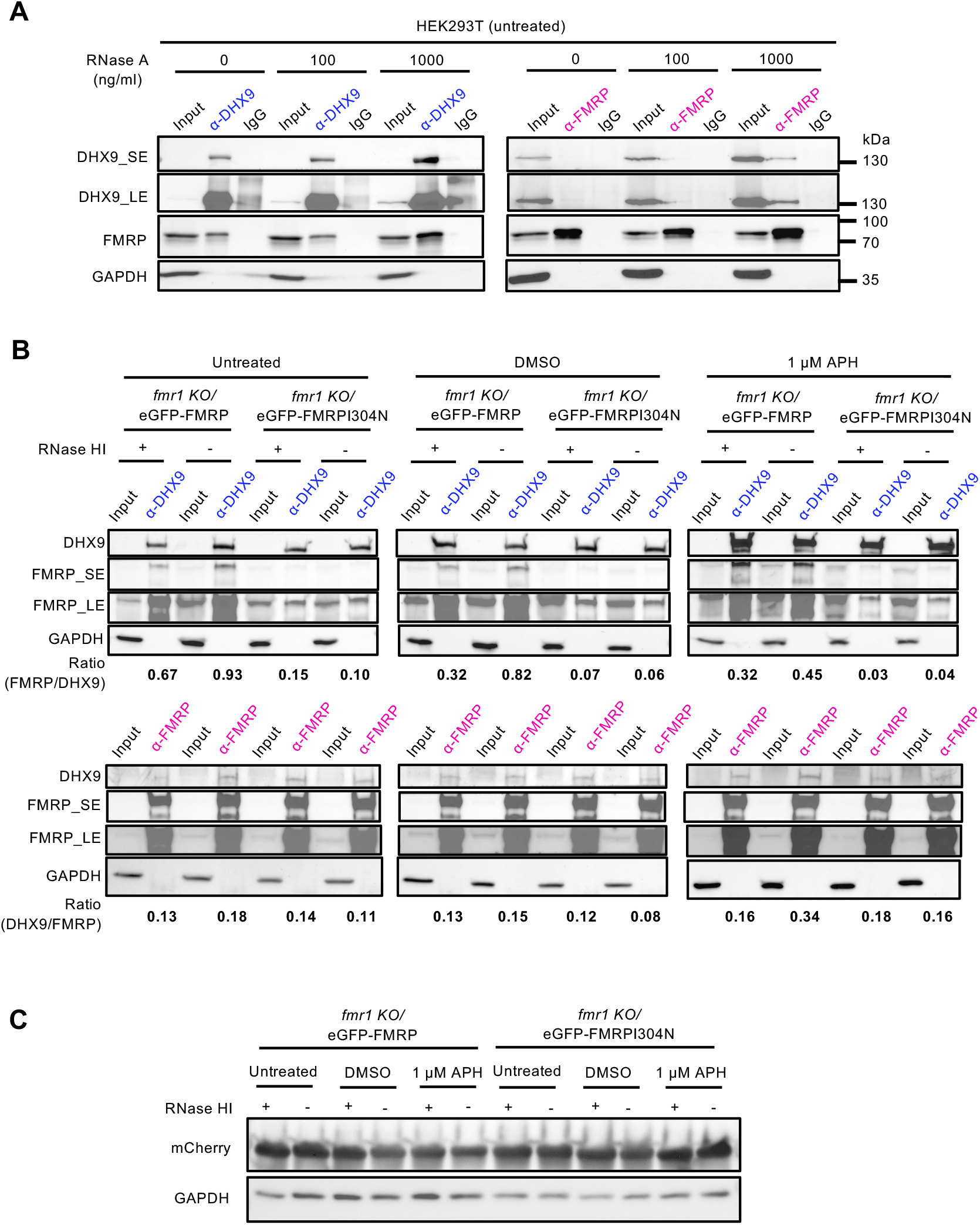
FMRP co-immunoprecipitates (IPs) with DHX9. **(A)** Co-IP of DHX9 and FMRP in HEK293T cells is not mediated by non-specific RNA. Increasing amounts of RNase A was added to the cell extract prior to IP reaction. (**B**) Co-IP of DHX9 and FMRP from *fmr1* KO cells expressing eGFP-tagged FMRP or FMRPI304N, and additionally, either WT RNase HI (“+”) or a catalytically dead RNase HI mutant (“-”). FMRP-I304N co-IPs less efficiently than FMRP. Cells were treated with DMSO, or 1 µM APH, or nothing. “SE”, short exposure. “LE”, long exposure. Numbers indicate ratio of Co-IP/IP. (**C**) Western blot showing similar expressions of WT RNase HI and catalytically dead RNase HI mutant tagged with mCherry in the indicated cell lines.

We performed an *in vitro* binding assay using recombinant histidine tagged-DHX9 (Figure S2E) and the aforementioned recombinant FMRP and fragments thereof. We added bezonase in the binding buffer to remove nucleic acids. Under these conditions, we indeed observed direct interaction between DHX9 and FMRP (Figure S4A). Moreover, this interaction specifically occurred through the N-Fold domain (Figure S4B). We note that the interaction was relatively weak, suggesting a transient/dynamic nature. Interestingly, the mutant N-Fold-I304N failed to interact with DHX9 (Figure S4C), indicating that the KH2 domain, an integral part of the N-Fold domain organization, is also important for recruiting R-loop resolution proteins. We then proceeded to test the hypothesis that FMRP recruits DHX9 to R-loops in the chromatin.

### FMRP regulates the chromatin association of DHX9

The above hypothesis predicts that i) DHX9 association with the chromatin increases upon APH treatment, and ii) DHX9 chromatin-association would be reduced/abolished in cells lacking FMRP or carrying the I304N mutation. To test these predictions we selected the β-actin locus previously shown to generate R-loops (Cristini et al., 2018), and analyzed protein association with sequences at the promoter, intron-5 and the pause site (for transcriptional termination) by chromatin immunoprecipitation coupled with quantitative PCR (ChIP-qPCR). We first analyzed chromatin binding by FMRP. With a noted exception at the promoter, FMRP-I304N showed similar level of association with the chromatin as the wild type FMRP (Figure 5A). The stronger association at the promoter by FMRP-I304N might reflect the higher affinity of the mutant for R-loop as shown by EMSA. We then analyzed DHX9 binding. To our surprise, in cells carrying the mutant FMRP-I304N (Figure 5B), or lacking FMRP entirely (Figure S5A), DHX9 showed increased presence at the promoter and pause sites, specifically during DMSO treatment. This observation suggested that DHX9 chromatin association was negatively correlated with the functional presence of FMRP, contrary to our original hypothesis. To test this hypothesis we asked if the subcellular localization of DHX9 was regulated by FMRP by comparing mCherry-DHX9 in control *FMR1^+/+^* cells and *fmr1KO* cells. Whereas mCherry-DHX9 showed pan staining pattern in the nucleoplasm in the control cells, it appeared to be reduced in the nucleoplasm and instead enriched in the nucleoli of the *fmr1KO* cells (Figure S5B). Moreover, complementation of the *fmr1KO* cells by expression of eGFP-FMRP partially reverted this phenotype (Figure S5B). Protein expression was confirmed by western blot (Figure S5C). These results supported our model in which FMRP regulates DHX9 chromatin binding dynamics.

**Figure 5.**
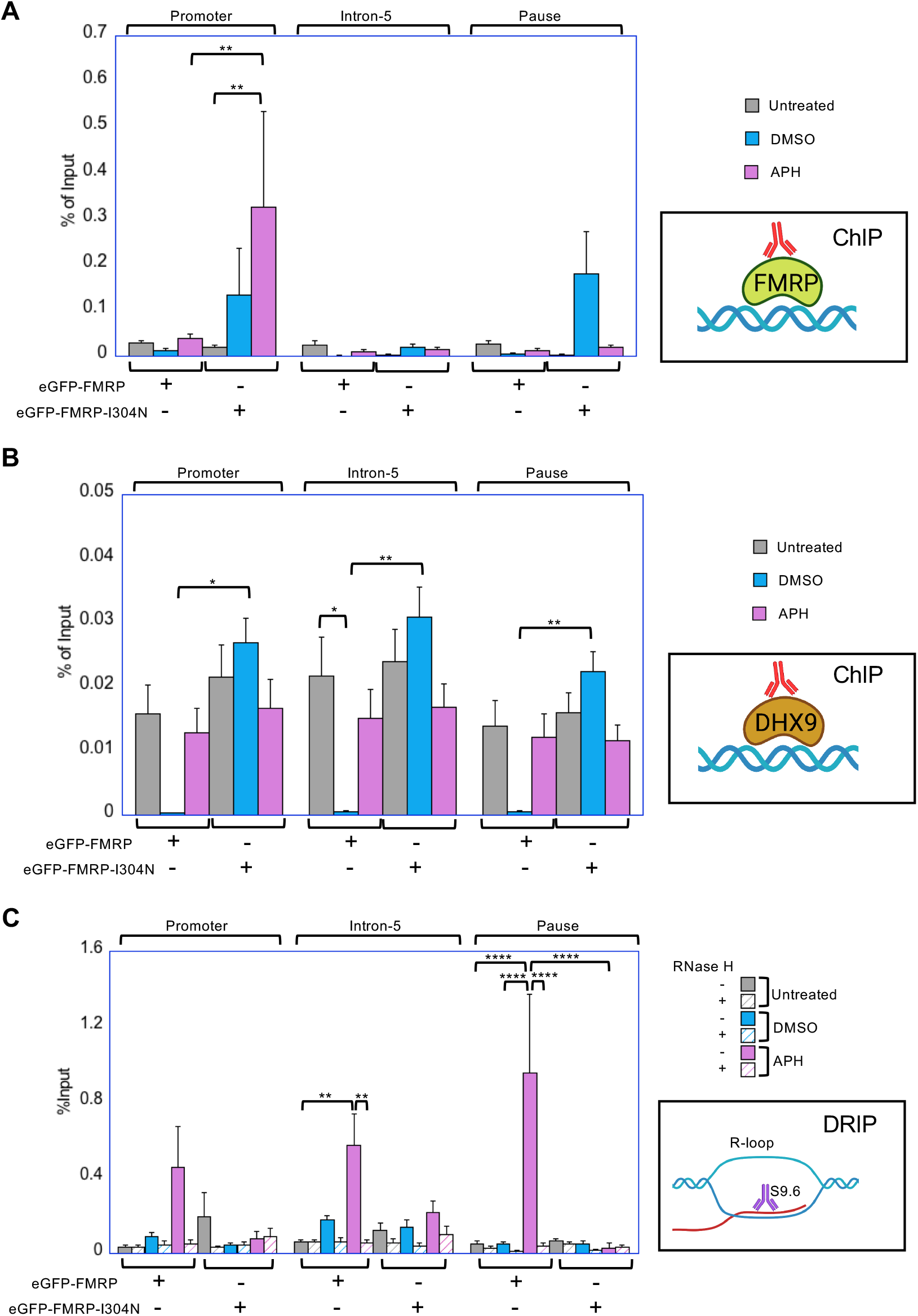
FMRP regulates chromatin association of DHX9. **(A)** FMRP ChIP-qPCR across the β-actin locus from *fmr1 KO* cells expressing eGFP-FMRP or GFP-FMRP-I304N. **(B)** DHX9 ChIP-qPCR across the β-actin locus from *fmr1 KO* cells expressing eGFP-FMRP, eGFP-FMRP-I304N. Two independent experiments were done and for each replicate and percentage of input was calculated. The average of the two experiments calculating percentage of input is shown for (A) and (B). **(C)** DRIP-qPCR across the β-actin locus from cells in (A). DRIP reaction was performed with and without RNase H treatment. Two independent experiments were done and average of the experiments are shown. Primers for the β-actin locus in promoter, intron-5 and pause sites were obtained from Cristini et al, 2018. One-way ANOVA and Sidak’s multiple comparison test were performed.

Meanwhile, we quantified R-loops by DNA:RNA immunoprecipitation (DRIP-qPCR) at the same loci which we observed DHX9 binding. Because the cells lacking FMRP and those carrying the FMRP-I304N mutant showed a similar phenotype of enhanced DHX9 chromatin association, we focused on the FMRP-I304N mutant hereon. In cells carrying wild type FMRP, DRIP signals increased with DMSO and APH treatment at all three loci, including the promoter, intron-5 and pause site, consistent with replication stress-induced R-loop formation (Figure 5B). Moreover, DRIP signals were sensitive to RNase H treatment, suggesting *bona fide* R-loop formation (Figure 5B). In comparison, the FMRP-I304N mutant cell line showed relatively lower DRIP-signals than FMRP across almost all conditions. Because we have confirmed that the mutant cells exhibited high level of DNA DSBs (Figure 4F) as well as R-loop formation based on αS9.6 staining (Figure 4G), we surmised that R-loop formation in the mutant led to DSBs and rendered the genomic substrate unamenable to DRIP detection as opposed to immunostaining by αS9.6. To test this hypothesis we analyzed the integrity of the genomic DNA at these loci by PCR. We predicted that the FMRP-I304N mutant cells would be deficient in DNA ampflification at the R-loop loci due to DNA breakage. We first confirmed that this result was not due to a deficiency in the genomic DNA in the mutant cells (Figure S5D). Indeed, we observed that PCR across the β-actin locus was significantly reduced in the mutant cells treated with APH (Figure S5E). Taking all these results together, there appeared to be an interesting dichotomy between FMRP and DHX9 where DHX9 chromatin association shows negative correlation with the functional presence of FMRP despite their physical interaction.

These results lent to a testable model in which we propose FMRP either blocks (physical sequestration) or disengages DHX9 from the R-loop template; however, we favor the latter possibility given the weak interaction between FMRP and DHX9. We speculated that DHX9, after unwinding the RNA:DNA duplex and upon reaching the 5’-junction of the R-loop, does not run off the trailing RNA spontaneously as the RNA might be tethered to other binding proteins, and instead requires a signal to disengage from the template. FMRP, by interacting with the three-way junction of the R-loop, may serve as a stop signal for DHX9 by transiently interacting with it and possibly curtailing its helicase activity, thereby disengaging it from the R-loop. We tested this model by first asking if FMRP alters the helicase activity of DHX9 *in vitro*.

### FMRP inhibits DHX9 helicase activity on R-loops through its N-Fold domain

DHX9 is a 3’ to 5’ helicase known to be able to unwind dsDNA, dsRNA as well as RNA:DNA hybrid, and in the context of RNA DHX9 demonstrated a clear preference for the 3’-RNA overhang (Dutta A et al., unpublished data and (Jain et al., 2010)). Therefore, we focused on an R-loop structure with a 3’-RNA overhang, which DHX9 readily unwound and produced free RNA (Figure 6A-C, lane 2 in all panels). When presented with increasing concentrations of FMRP-WT the DHX9 helicase activity was steadily reduced, reaching near complete inhibition at 400 nM (Figure 6A). The inhibition was more pronounced when N-Fold-WT was added at the same concentrations (Figure 6B). The I304N mutation in both the full length and N-Fold contexts reduced the inhibitory effect (Figure 6A&B). Finally, the C-IDR did not appear to inhibit DHX9 (Figure 6C). Quantification of % R-loop unwinding from three independent experiments confirmed these observations (Figure 6D). Therefore, we concluded that FMRP inhibits DHX9 via its transient interaction with DHX9 and not simply through competing for R-loop. These characteristics of the negative impact on DHX9 by FMRP during R-loop unwinding are consistent with our model.

**Figure 6.**
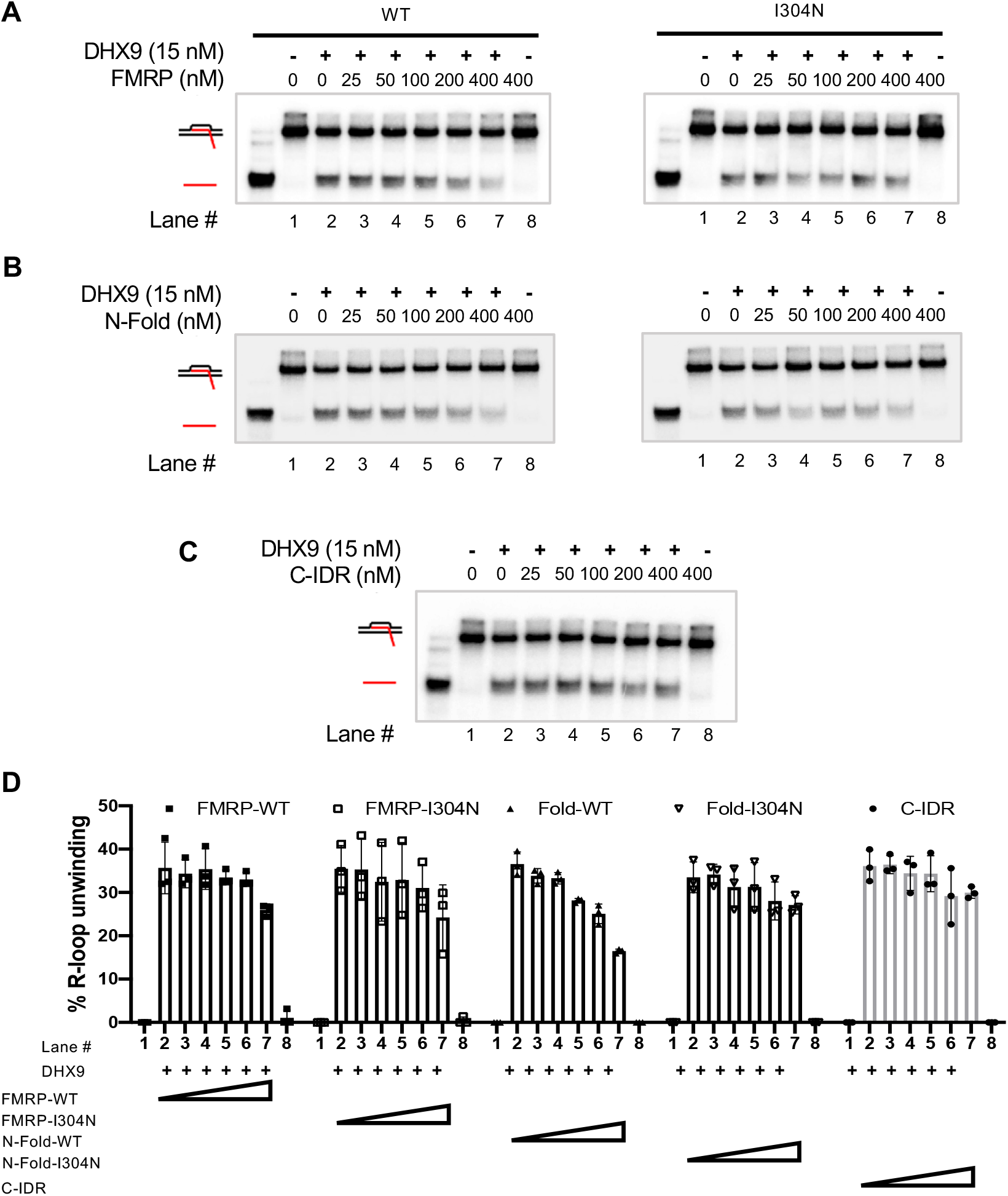
FMRP regulates DHX9 helicase activity. **(A)** High concentrations of FMRP full length WT and I304N inhibits the helicase activity of DHX9 on R-loop with 3’ overhang *in vitro.* **(B)** N-Fold WT inhibits DHX9 helicase activity more than I304N. **(C)** C-IDR does not affect DHX9 helicase activity. Black strand represents DNA and red represents RNA. **(D)** Quantification of DHX9 helicase activity shown here as percentage R-loop unwinding for full length, N-Fold of both WT and mutant FMRP and C-IDR domain.

## DISCUSSION

The work described here is directly predicated on our recent study demonstrating a genome protective role of FMRP in preventing replication stress-induced R loop formation and DSBs (Chakraborty et al., 2020). Here we provided additional support for this novel function of FMRP by demonstrating in a CRISPR KO of *fmr1* model that DNA damage and R-loop formation are both elevated. We further demonstrated association between FMRP and R-loop both in vivo and in vitro. We showed, for the first time, that the C-IDR of FMRP can interact tightly with the R-loop structure. This is a remarkable finding, given that the same C-IDR also has the ability to interact with G-quadruplexes and SoSLIPs that both adopt very different 3D structures than R-loops (Bechara et al., 2009; Santoro et al., 2012; Vasilyev et al., 2015). Previous studies have demonstrated that the formation of R-loops and G-quadruplexes are potentially coupled during transcription (De Magis et al., 2019; Lee et al., 2020). Together with our finding, it appears that FMRP can bind both structures via its C-IDR, thus providing a mechanism for the functional linkage between these non-canonical nucleic acid strutures.

Based on the hierarchy of substrate binding by the FMRP segments, we propose that upon replication stress FMRP binds to R-loops predominantly via its C-IDR, thereby allowing the KH domains to bind the trailing nascent ssRNA, and the N-terminal Agenet domains to presumably interact with methylated histone tails or R-loop resolving factors that contain motifs with methylated arginine or lysine residues. Here, we showed that FMRP interacts with one such R-loop resolving factors, DHX9, through its N-Fold domain. Moreover, the interaction is dependent on a bona fide KH2 domain, suggesting that mutations in the KH domain may interfere with the Agenet domain’s binding to other proteins through disruptions of intra-molecular interactions. Furthermore, our *in vitro* experiments suggested that in the absence of other proteins, the N-Fold domain can adopt intramolecular interaction with the C-IDR domain thereby interfering with the C-IDR binding to R-loop. Interestingly, the I304N mutation apparently reduced such interference, while it also reduced the abilit of the N-Fold to interact with DHX9 or R-loop. These results underscore the importance of the KH2 domain in proper FMRP function, and the direct impact on disease manifestation. Future experiments will be directed towards understanding the mechanism governing such intramolecular interactions--for instance--whether/how posttranslational modifications might play a role in such a mechanism.

Unexpectedly, we found that FMRP inhibits the DHX9 helicase activity on R-loops *in vitro*. Moreover, the chromatin association of DHX9 was lower in the presence of a functional FMRP than in its absence. These results led us to propose the following “disengagement” model to describe the complex interplay between FMRP and DHX9 at R-loops (Figure 7). We propose that FMRP and DHX9, each capable of binding R-loops directly, are associated with the 5’-and 3’-end of the R-loop, respectively. This mode of interaction is based on the observed substrate preference of FMRP and the known preference of DHX9 for substrates with a 3’-overhang and its association with RNA Pol II. Once engaged on the 3’end of the RNA DHX9 unwinds the DNA:RNA hybrid within the R-loop towards the 5’ end. We note that very little is known about how DNA translocases disengage from the *in vivo* substrates, a problem that likely does not exist in *in vitro* assays as the substrate has finite length and is not tethered to the chromatin. Therefore, we propose that once poised at the 5’ end three-way junction of the R-loop, FMRP functions as a signal for DHX9 to stop translocation and disengage from the chromatin upon completion of unwinding the DNA:RNA hybrid. Importantly, such a function would require the interaction between the two proteins relatively weak, consistent with our observation. In the absence of a functional FMRP such as the FMRP-I304N mutant, FMRP fails to eject DHX9 and allows the RNA strand to re-enter to form the R-loop, ultimately causing DSBs. Consistent with this model we have also observed that FMRP regulates the subcellular localization pattern of DHX9.

**Figure 7.**
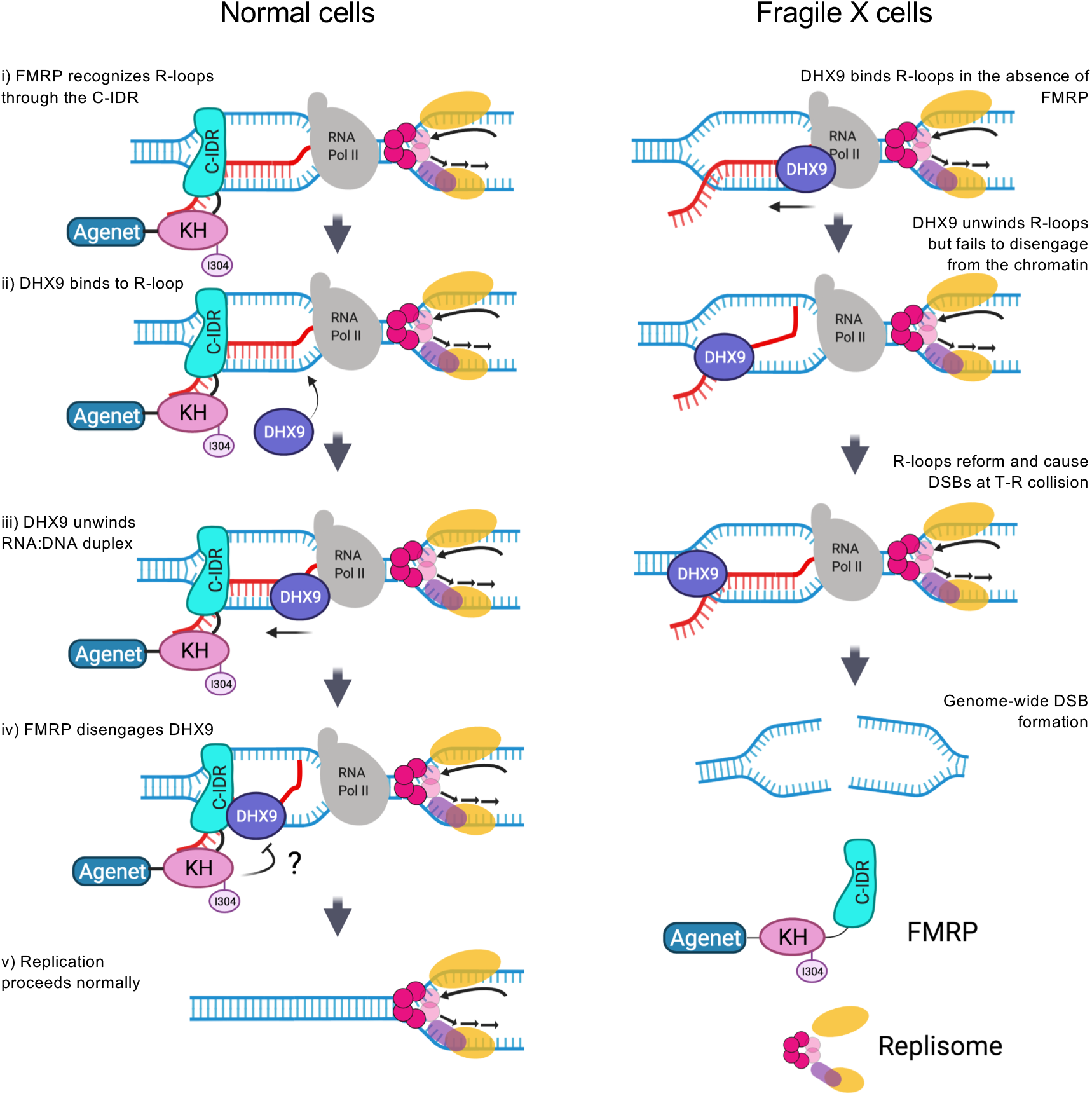
Proposed model for the functional interplay between FMRP and DHX9 at the chromatin.

However, we note that a competing “blockage” model must also be entertained (not depicted). It has been suggested that in certain genetic background such as splicing mutations, DHX9 promotes R-loop formation by unwinding the secondary structure in the RNA transcript and permitting its re-entry into the DNA template to form hybrids (Chakraborty and Grosse, 2011). It is possible that FMRP actively curtails the helicase activity of DHX9 on dsRNA and thereby prevents R-loop formation. We have used an *in vitro* assay with radioactively labeled dsRNA and cold DNA bubble to measure R-loop formation after DHX9 unwinds the dsRNA to allow it to enter the DNA bubble. Thus far, we have not observed any evidence of FMRP having an impact on this activity of DHX9 (data not shown). However, we note that the *in vitro* dsRNA substrate is not a preferred substrate for FMRP, which might mask the potential impact by FMRP on the DHX9 helicase activity on dsRNA. Additionally, when evaluating the impact of this “disengagement” model for the DHX9-FMRP interplay we must also consider the fact that FMRP interacts with numerous proteins, though predominantly cytoplasmic proteins. It is conceivable that FMRP also regulates other R-loop enzymes, where it assumes a different role than it does in the context of DHX9. A recent human interactome analysis in HeLa cells revealed an interaction between FMRP and the THO-TREX complex, which functions at the interface of transcription elongation and mRNA export (Hein et al., 2015). THOC1, a subunit of the THO/TREX complex was present in the same complex as FMR1, DHX9 and other THOC proteins. Depletion of subunits in the hTHO complex causes DNA damage that is R-loop dependent (Dominguez-Sanchez et al., 2011). Our co-immunoprecipitation experiments also showed an interaction between FMRP and TOP3B, whose loss causes R-loop-mediated genome instabilty (Zhang et al., 2019). This result suggests that FMRP forms multiple docking sites for factors that resolve R-loops and ensures proper transcription, RNA processing and export.

Finally as a closing throught, modular proteins such as FMRP and DHX9, which contain multiple folded domains interspersed with intrinsically disordered regions, often undergo liquid-liquid phase separation (LLPS), where molecules spontaneously demix from their solvent to form their own microscopic droplets (Banani et al., 2017; Forman-Kay et al., 2018; Holehouse and Pappu, 2018). The C-IDR of FMRP is capable of undergoing LLPS in isolation, in the context of full length, and in the presence of its cognate RNA substrates (Tsang et al., 2019). The multivalent interactions with diverging K_D_s between various FMRP segments, R-loop substructures and R-loop resolving factors (*e.g.*, DHX9) can be the basis for the assembly of a phase-separated, membrane-less foci for resolving R-loops (Dettori et al., 2021).

## ACKNOWLEDGEMENT

We thank Drs. Leszek Kotula, Frank Middleton, Patricia Kane for helpful discussions. We thank Dr. Helmut Pospiech and Late Dr. Frank Große from Leibniz Institute for Age Research -Fritz Lipmann Institute (FLI) at Jena, Germany for providing P. Sung with the pFastBac-His-DHX9 construct. We also thank Charlotte Logan for assisting in cloning of FMRP mutant constructs and Nathan McKean for helping with protein purification of FMRP. R-loop structures in Figure 2 and model in Figure 7 were created with BioRender.com.

This work was supported by the National Institute of Health GM118799-01A1 grants to W.F., the Department of Defense CDMRP Discovery award W81XWH-15-1-0204 to W.F., the Department of Defense grant PC160083 to H.H., the National Institute of Health grant R35CA241801 to P.S. and the National Institute of Health Award R35GM138097 to A.B..

## AUTHOR CONTRIBUTION

A.C. and W.F. conceived the study. A.C. performed and analyzed the immunostaining experiments with help from H.H. A.C. performed chromatin fractionation, co-immunoprecipitation, ChIP-qPCR and DRIP-qPCR experiments with guidance from W.F. A.C. generated HEK293T *FMR1* CRISPR KO clones and EGFP-FMRP WT or mutant clones from pooled cells. J.L. generated EGFP-FMRP, EGFP-FMRPI304N pool and mCherry-DHX9 and mCherry-DHX9-HD expressing pool of stable cell lines. JL performed IP for mCherry-DHX9 expressing cells. L.D. and A.B. designed and performed protein purification for all full length FMRP, FMRPI304N and FMRP domains. L.G. and X.X. performed some preliminary EMSA assays. A.D. designed and performed EMSA assays, purified DHX9 and performed FMRP:DHX9 interaction assay, with P.S. providing experimental guidance and suggestions. A.C., A.B. and W.F. wrote the manuscript with input from all authors.

## MATERIALS AVAILABILITY STATEMENT

This study generated a collection of plasmids, cell lines and recombinant proteins. All materials will be distributed upon request after publication.

## MATERIALS AND METHODS

### Cell line growth and culture conditions

Human EBV transformed lymphoblastoid cell lines, GM06990 (control) and GM03200 (Fragile X) were obtained from Corielle institute. Lymphoblastoids were grown in RPMI1640 (Corning), supplemented with GlutaMAX (GIBCO), 15% heat-inactivated FBS (Fetal Bovine Serum, Benchmark), 100 IU/mL penicillin and 100 μg/mL streptomycin (Corning) at 37°C with 5% CO_2_. HEK293T (ATCC #ACS-4500) cells and Phoenix-AMPHO producer cells (ATCC #CRL-3213) were grown in DMEM medium (GIBCO) supplemented with 10% FBS, 1X GlutaMAX, 100 IU/mL penicillin and 100 μg/mL streptomycin, 1 mM sodium pyruvate (Corning), 10mM HEPES buffer (Corning) and 1X MEM non-essential amino acids (Corning) and grown at 37°C with 5% CO_2_.

### Generation of CRISPR KO of FMR1

FMR1 sgRNA CRISPR/Cas9 All-in-One Non-viral Vector set (Human) (abm # K0790727) containing three targets (T1,T2 and T3) were used to create FMR1 knock-out lines in HEK293T cells. Additionally, CRISPR Scrambled sgRNA All-in-One Non-Viral Vector (with spCas9) (abm #K094) was used as control for the knock-out. HEK293T cells were seeded onto 60 mm plates at 80% confluency and transfected using DNAfectin Plus Transfection reagent (abm) following manufacturer’s instruction. Briefly, 5 μg of each construct (Scr, T1, T2 and T3) was mixed with serum-free and antibiotic free media. 15μl of the transfection reagent was added to this mixture and incubated for 30 m in room temperature. Following incubation, the mixture was added drop-wise to the cells after 20 h of seeding. 48 hr post transfection, cells were trypsinized, washed and resuspended in FACS buffer (2% FBS in 1X PBS) and filtered through filter-topped flow tubes (BD falcon) using a luer-lock syringe at 2×10^6^ cells/ml. Cells were sorted and selected for mid-intensity GFP signal using untransfected cells as a control. Single cells were seeded on to 96-well plate containing media for generating clones for all targets and the scramble. 10-11 clones per target and scramble were further expanded for western blot analysis of FMRP expression. Ultimately, 3-5 clones per target showing optimal loss of FMRP expression (no visible FMRP expression) was selected for further analysis. Genomic DNA was isolated from these clones using CRISPR genomic cleavage detection kit (abm) and PCR amplification of sequences around the target region using primers; FMR1_T1_F: 5’-CTCAGCTCCGTTTCGGTTT-3’, FMR1_T1_R: 5’-AAAGGGGGAATAAGCCATCG-3’, FMR1_T2_F: 5’-ATTGCCGTTATGTCCCACTC-3’, FMR1_T2_R: 5’-TCAACGGGAGATAAGCAGGT-3’, FMR1_T3_F: 5’-CTGCCTACCTCGGGGTACAT-3’, FMR1_T3_R: 5’-GCTCTTGCAAACCAAACCAT-3’, was conducted. The PCR product was then sequenced and the sequences were analyzed to verify substitution, addition or deletion of nucleotides at the target region indicating a mutation and leading to loss of FMRP expression. B3 clone of Target-3 was used for the rest of the experiments and for generation of EGFP alone, EGFP-FMRP and EGFP-FMRPI304N fusion protein expressing stable cell lines.

### Generation of EGFP-fusion protein stable cell lines

Plasmids expressing FMRP and FMRPI304N was generated as described previously (Chakraborty et al., 2020). mCherry was PCR amplified from mCherry-Alpha-5-Integrin-12 (Addgene #54970) using forward primer 54970RMmCherryaddNHis_F2 : 5’-CGAGGTTAACATGGGCCATCATCATCATCATCATATGGTGAGCAA-3’, and reverse primer, 54970RMmcherry_R1: 5’-CCATGAATTCCTTGTACAGCTCGTCCATGCCGCCG-3’, cloned into pMSCVpuro (Addgene #K1062-1) at Hpa1 and EcoR1 sites to create pMSCVpuro-His-mCherry. DHX9 and DHX9 helicase dead mutant (DHX9-HD) were PCR amplified from pFBDual-DHX9 and pFBDual-DHX9 helicase dead mutant (gifts from Sung lab), using forward primer pFB_rmdhx9+N3AAs_F2 : 5’-ACCCGAATTCAACTTGGTTATGGGTGACGTTAAAAATTTTCTG-3’, and reverse primer, pFB_RMDHX9_R1: 5’-GGTAGAATTCTTAATAGCCGCCACCTCCTCTTCC-3’, and cloned to pMSCVpuro-His-mCherry at EcoR1 site to create pMSCVpuro-His-mCherry-DHX9 and pMSCVpuro-His-mCherry-DHX9-HD. These constructs were then packaged into retrovirus using Phoenix-AMPHO producer cells (ATCC #CRL-3213) as described previously (Chakraborty et al., 2020). Viral particles so generated were then used to transduce *fmr1* KO-clone B3 or Scramble cells and generate pooled population of EGFP-FMRP, EGFP-FMRPI304N, EGFP alone and maintained in 0.25 μg/ml puromycin containing media. Cells were sorted and selected for mid-intensity GFP signal using the parent *fmr1* KO-clone B3 cells as a control. EGFP expressing single cells were seeded on to 96-well plate containing media for generating clones for all cell lines. Pooled cells expressing both EGFP and mCherry signals were selected with FACS. Expression of respective proteins were verified through microscopy and western blot.

### Co-immunoprecipitation (Co-IP)

Approximately 6-7×10^6^ cells were used for each IP reaction. Cells were resuspended in 1 ml IP lysis buffer [25 mM Tris-HCl pH 7.5 / 150 mM NaCl / 1% NP-40 / 1 mM EDTA / 5% glycerol / Halt protease inhibitor cocktail (Thermo scientific) / Halt phosphatase inhibitor cocktail (Thermo scientific)] and incubated on ice for 1 h. Cell lysates were sonicated to fragment the chromatin and reduce viscosity followed by centrifuged at 10,000 rpm for 10 m. Protein concentration in the supernatant was determined using Pierce protein assay reagent (Thermo Scientific). Fifty micro-liter of Dynabeads protein G (Invitrogen) per reaction were incubated with 200 μl antibody binding buffer [1X PBS/ 0.02% Tween 20] and 5 μg of anti-FMRP (Biolegend) and anti-FXR1(Santa Cruz Biotechnology #sc-374148), or 4 μg anti-DHX9 (Santa Cruz Biotechnology #sc-137232), 5 μg mouse IgG control (Biolegend) or 4 μg rabbit IgG (Bethyl laboratories) in a rotator for 10 m at room temperature. The immuno-complex was rinsed with 200 μl antibody binding buffer at room temperature, followed by incubation with 500 μg of cell lysate per reaction at 4°C overnight. After incubation the supernatant was saved as flow-through (FT) and the beads were washed twice with IP lysis buffer without NP-40. 50 μl 2X Laemmli buffer was added to the beads and boiled for elution, before analysis on 8% SDS-PAGE or gradient (4-15%, BioRad) gels and western blotting using anti-FMRP (Cell signaling #LS-C82231, 1:500 or Biolegend #6B8, 1:1000), anti-GAPDH (Santa Cruz Biotechnology #sc-47724, 1:4000) or anti-DHX9 (Santa Cruz Biotechnology #sc-137232, 1:500), anti-Top3β (Santa Cruz Biotechnology #sc-137238, 1:1000) anti-FXR1 (Santa Cruz Biotechnology, 1:1000), anti-GFP (Santa Cruz Biotechnology #sc-9996, 1:1000) and anti-mCherry (Santa Cruz Biotechnology #sc-390909, 1:100). For RNase A treatment experiments, lysates were first divided into equal protein aliquots and treated with multiple concentrations (as shown in figure) of RNase A or left untreated for 20 minutes on ice. These lysates were then used for IP as described above.

### RNase H I overexpression

pICE-RNaseHI-NLS-mCherry (Addgene #60365) and pICE-RNaseHI-D10R-E48R-NLS-mCherry (Addgene #60367) was transfected into FMR1KO-B3-FMRP-H9 and FMR1KO-B3-FMRPI304N-F10 cells using TransIT-2020 transfection reagent (Mirius Bio) following manufacturer’s instruction. Twenty-four hours after transfection, cells were treated with DMSO and APH separately or left untreated. Forty-eight hours after transfection lysates were prepared for co-immunoprecipation as described above.

### Subcellular fractionation

Cells were grown to a density of 0.4-0.5×10^6^ cells/ml with >90% viability. Cells were treated for 24 h with aphidicolin, DMSO or nothing. Samples were collected as aliquots of approximately 5×10^6^ cells, washed twice with PBS, then frozen for storage. Each thawed aliquot of cells was resuspended in 500 μl Farham’s lysis buffer without NP-40 [5 mM PIPES pH 8.0 / 85 mM KCl / Halt protease inhibitor cocktail] and incubated on ice for 2 m. 50 μl of the cell lysate thus prepared was collected as a whole cell extract control and the remaining lysate was spun at 1300 g for 4 m to pellet nuclei. The supernatant served as the crude cytoplasmic fraction. The nuclear pellet was resuspended in 150 μl Farham’s lysis buffer and incubated for 20-30 m at 4°C and served as the nuclear fraction. Equal volume of 2X Laemmli buffer were added and samples were boiled and later sonicated. Approximately 3×10^5^ cell equivalent per fraction was used for electrophoresis on a 12% SDS-PAGE gel, followed by western analysis. Densitometry of autoradiogram was done using ImageJ (https://imagej.nih.gov/ij/) to calculate the percentages of FMRP in the nuclear and cytoplasmic fractions.

#### Western blot

Whole cell lysates were prepared in lysis buffer [50 mM Tris-HCl pH 7.5 / 0.5 M NaCl / 10 mM MgCl_2_ / 1% NP-40 / Halt protease inhibitor cocktail / Halt phosphatase inhibitor cocktail] and at least 20 μg of proteins were analyzed by 10% SDS-PAGE before western blotting. The following antibodies were used: anti-FMRP (Biolegend, 1:1000), anti-Histone H3 (Cell Signaling, 1:500) and anti-GAPDH (Santa Cruz Biotechnology, 1:2000).

### Immunocytochemistry and microscopy

#### Lymphoblastoid cells

Approximately 4×10^5^ lymphoblastoid cells after 24 h drug treatment described above were pelleted, washed and resuspended in 500 ml 1X PBS. Cells were seeded onto 0.5 ug/ml poly-D-lysine (Sigma-Aldrich) coated coverslips in a 24-well plate. Cells were allowed to adhere for 5 m. 125ml of 4% paraformaldehyde was used to spike the cells for 2 m at room temperature, followed by removal of the solution and replaced by fresh 500 ml 4% paraformaldehyde. Cells were washed three times with 1XPBS and permeabilized for 30 m with permeabilization buffer (0.5% Triton-X in 1XPBS). Following permeabilization, cells were subjected to RNase H (15U/well) or RNase H (NEB) buffer only treatment for 4 hr and RNase III or RNase III buffer (Ambion, Invitrogen) only for 30 m at 37°C. Enzyme treatment was followed with two washes with 1X PBS. *HEK293T cells*: 5×10^4^ cells were seeded onto 0.1 ug/ml coated poly-D-lysine coverslips in a 24-well plate and cultured for 36 h. At 70 % confluency, cells were treated with drugs for 24 hr. Post treatment, cells were fixed with 500 μl 2% paraformaldehyde for 20 m at room temperature followed by gentle washing with PBS three times. Cells were then blocked with 500 μl PBSAT (1% BSA, 0.5% Triton X in PBS) for 1 h at room temperature. Fifty microliter of primary antibody solution was applied to all coverslips and incubated overnight at 4°C, washed with PBSAT, and incubated with 50 ml secondary antibody for 1 hr at room temperature. Cells were then washed with PBSAT followed by PBS and mounted on glass slides using mounting media (Prolong Diamond antifade plus DAPI, Invitrogen). Coverslips were allowed to solidify for 24 h before imaging on Leica SP8 confocal fluorescence microscope. Antibodies used for immunostaining include the following: primary antibodies: anti-γH2A.X, Cell Signaling #9718S, 1:400, anti-FMRP, Cell signaling/Biolegend, 1:200; S9.6, Kerafast #ENH001, 1:250; anti-LaminA+C, Novus Biologicals # NBP2-25152, 1:500, and secondary antibodies: Alexa fluor 488, 568, and 647, Invitrogen # A-21206, A10037 and A-21449, respectively, 1:400. To determine localization of FMRP and R-loop in the nucleus, single plane images were obtained. For measurement of S9.6 signal, a region of interest (ROI) in unperturbed images of DAPI was used, which was overlayed on S9.6 signal and Fiji (https://imagej.net/software/fiji/) was used to measure integrated density of the ROI. For measuring colocalization, Coloc2 plugin in Fiji was used with DAPI as ROI. Manders’s overlap co-efficient for calculated for both the channels and tM1; FMRP’s overlap with RNA:DNA hybrids was used to calculate percentage overlap as shown in Figure 1D and S1E. For the purpose of presentation, images were adjusted for background and contrast and smoothed using a gaussian blur of 1 in Fiji and representative images were used (identical adjustments have been made for FMRP and S9.6 signals for all samples in Figure 1 and Supplementary Figure 1). To quantify DNA damage, γH2A.X signal in nucleus was measured from single plane images. A region of interest (ROI) in unperturbed images of DAPI was used and overlayed on γH2A.X signal. Fiji was used to measure integrated density of the ROI.

#### Live-Cell Experiments and Imaging

Stably transfected cells co-expressing EGFP-FMRP and mCherry-DHX9, or EGFP-FMRP and mCherry-DHX9-HD were grown at 37 °C (5% CO_2_) in DMEM (Gibco) supplemented with 10% FBS (Benchmark). Cells were plated and cultured in 12 Well glass bottom plates (Cellvis) before live-cell imaging. Images were taken on a Leica stimulated emission depletion (STED) 3X nanoscope with a 93X glycerol objective.

### Cloning and protein purification

As previously outlined in Tsang et al (Tsang et al., 2019) and briefly described here, codon optimized full length human FMRP Isoform 1 cDNA was generated by gene synthesis (GeneScript, Inc) and was subcloned into a pET-SUMO vector (Invitrogen). This pET-SUMO-FMRP plasmid was used as a template to generate (i) full length I304N mutant, (ii) FMRP-WT and FMRP-I304N mutant N-Folds (residues 1-455 without and with the I304N substitution, respectively), and (iii) C-IDR (residues 445-632) via QuikChange Site-Directed Mutagenesis (Agilent) for protein expression. The fidelity of these constructs was confirmed by Sanger sequencing (Eurofins Genomics, Louisville, KY). Each construct was transformed into *Escherichia coli* BL21(DE3) Codon Plus Cells (Agilent). Select colonies were inoculated in 50 ml of Luria Broth (LB) medium, before dilution into 1 L fresh LB medium in a Fernbach flask and grown at 37°C. Protein expression was induced with 1 mM isopropyl-β-D-thiogalactopyranoside (IPTG) at an optical density (600 nm) of ∼0.6 and was incubated at 16°C for 18 h. Cells were harvested by centrifugation at 15,000 rpm for 30 m. The supernatant was carefully discarded, and each cell pellet was stored at −20°C until ready for protein purification.

To begin purification, frozen cell pellets were thawed and re-suspended in 100 ml of lysis buffer containing 100 mM NaCl, 50 mM Na_2_PO_4_, 200 mM Arginine HCl, 200 mM Glutamic acid, 10% Glycerol, 10 mM β-mercaptoethanol, and 1% CHAPS, pH 7.4, supplemented with DNase I, lysozyme and protease inhibitors (bestatin, pepstatin, and leupeptin). Cells were lysed by sonication and the lysate was subjected to centrifugation at 15,000 rpm for 30 m. The supernatant was loaded onto a 20 ml HisTrap HP column (GE Healthcare) equilibrated in the binding buffer (i.e. same composition as lysis buffer, but without DNase I and lysozyme) and incubated at 4°C for 30 m. The column was extensively washed three times with 30 ml of the equilibration buffer. SUMO-fusion proteins were eluted using the same equilibration buffer supplemented with 500 mM imidazole, and fractions containing proteins were combined. A 6X-His-tagged Ulp protease was added to cleave the His-SUMO tag at room temperature overnight with rocking. Completion of the Ulp cleavage reaction was confirmed by SDS-PAGE. After cleavage, the protein solution was passed through a 0.2 µm filter to remove any aggregated product, before it was concentrated using a 5 kDa-cutoff Amicon concentrator by centrifugation at 4,000 rpm at room temperature. The concentrated protein solution is again filtered before being loaded onto an equilibrated Superdex 200 size exclusion column (GE Healthcare) to separate the FMRP constructs from the Ulp protease and the His-SUMO fusion tag. Fractions containing pure FMRP proteins were identified by SDS-PAGE and combined for storage at −80°C.

DHX9-His was expressed by transducing 800 ml Tni cell culture in ESF921 serum-free media (Expression Systems) at a density of 1×10^6^ cells/ml with 16 ml baculoviral suspension (generated in Sf9 cells) and grown for 70 h at 27°C with shaking. Cell pellet was resuspended in a lysis buffer containing 50 mM Tris-HCl, pH 7.5, 500 mM KCl, 10% glycerol, 1 mM EDTA, 1 mM DTT, 0.01% NP-40, 2 mM ATP, 4 mM MgCl_2_, 10 mM Imidazole, cOmplete protease inhibitor cocktail (MilliporeSigma), and 1mM PMSF, with sonication. The lysate was clarified by ultracentrifugation at 40,000 rpm for 45 m. The clarified lysate was incubated with 1 ml Ni-NTA resin (Qiagen) for 1 h, followed by washing the resin with 400 ml wash buffer-A containing 50 mM Tris-HCl, pH 7.5, 1000 mM KCl, 10% glycerol, 1 mM EDTA, 1 mM DTT, 0.01% NP-40, 4 mM ATP, 8 mM MgCl_2_ and 20 mM Imidazole. Protein-bound resin was washed again with 50 ml wash buffer-B containing 50 mM Tris-HCl, pH 7.5, 100 mM KCl, 10% glycerol, 1 mM EDTA, 1 mM DTT, 0.01% NP-40, and 20 mM Imidazole, followed by elution with 10 ml elution buffer containing 50 mM Tris-HCl, pH 7.5, 100 mM KCl, 10% glycerol, 1 mM EDTA, 1 mM DTT, 0.01% NP-40, 300 mM Imidazole and cOmplete protease inhibitor cocktail (MilliporeSigma). The elution was subjected to ion exchange purification with equilibrated Hitrap SP HP (1 ml) column at a gradient of 100-500 mM KCl. The peak fractions containing the protein were pooled together and purified again with HitrapQ (1 ml) column. The peak fraction was aliquoted, flash frozen with liquid nitrogen and stored at −80°C. The protein was also evaluated via size exclusion chromatography by loading 400 µl of the Hitrap SP HP purified fraction onto Superdex 200 increase 10/300 GL column (GE Healthcare), and a monodisperse peak was obtained at 11.8 ml elution fraction.

### Electrophoretic mobility shift assay (EMSA)

DNA or RNA was labeled at 5′-termini with T4-Polynucleotide kinase (NEB) using γ-P^32^-ATP as indicated in Figure 2. The oligo sequences are listed in Table S1. R-loops, RNA-DNA hybrids or duplex DNA substrates were generated by annealing the labeled oligonucleotide with the complementary cold oligonucleotides in equimolar ratio, as indicated in Table S2, by gradually decreasing temperature from 95°C to 4°C. Prior to binding assays all the substrates were checked by electrophoresis in 5% native TAE (30 mM Tris-acetate, pH 7.4 and 0.5 mM EDTA) polyacrylamide gel.

#### Binding assay

1 nM of R-loop, RNA-DNA hybrid, dsDNA, bubble DNA, ssDNA, or RNA substrate was mixed with 1 µl of protein at concentrations indicated in Figure 2, in a buffer composed of 25 mM Tris-HCl (pH 7.5), 100 mM KCl, 5 mg/ml BSA, 5 mM EDTA, with a final volume of 10 µl. This mixture was incubated 30 m on ice, followed by addition of 2 µl loading buffer composed of 50% glycerol, 20 mM Tris-HCl, pH 7.4, 0.5 mM EDTA, 0.05% Orange G.

#### Polyacrylamide gel electrophoresis

Electrophoretic separation of the protein-bound substrates was carried out by running the mix in 5% native TAE gels, at 110V for 90 m at 4°C. The gels were vacuum dried for 30 m at 80°C on a gel dryer and exposed to phosphorimaging screen overnight. Imaging was done using Typhoon molecular imager (Amersham) and bands were quantified using ImageQuant TL 8.0 image analysis software.

#### Image analysis

Images obtained were then used to perform band intensity analysis. Intensity of non-shifted and shifted band was measured. After a background correction, percentage of band shift was calculated for at least two replicates. The average of the replicates was then used to plot a scatter plot followed by nonlinear regression (curve fit) using Prism version 9. We used ‘specific binding with hill slope’ analysis for full length proteins to calculate dissociation constants (K_D_) and ‘one-site-Total’ for N-fold and C-IDR domains listed in Table 1.

#### *In vitro* protein binding assay (for FMRP protein domains and DHX9-His)

5 µg DHX9-His was incubated with 10 µl Ni-NTA beads in a binding buffer containing 50 mM Tris-HCl, pH 7.5, 150 mM KCl, 10% glycerol, 1 mM EDTA, 1 mM DTT, 0.01% NP-40, 0.1% Tween-20, 10 mM Imidazole, and 1 µl benzonase (MilliporeSigma) for 1 h, with mild shaking at 4°C. The supernatant was removed, and beads were washed three times with 200 µl binding buffer. The binding buffer was completely removed and DHX9-His bound Ni-NTA were further incubated for 15 m with 5 mg FMRP (full length)-WT, N-Fold-WT, N-Fold-I304N, or C-IDR (as indicated in the figures) in 20 µl binding buffer. The protein bound resins were spun down and the supernatants were taken out carefully. 5 µl loading buffer was added to supernatants. The resins in each tube was washed three times with 200 µl wash buffer (same buffer with 20 mM Imidazole, and 200 mM KCl, without benzonase). The bound proteins were eluted with 25 µl 1X Laemmli buffer. Equal volume of supernatants and the pulldowns were analyzed in 4-15% polyacrylamide gradient gel.

### R-loop unwinding assay

2 nM R-loop with 3’-RNA overhang (5’-γP^32^ labelled) was incubated with 15 nM DHX9 for 30 m at 37°C in a buffer consisting 10 mM Tris-HCl, pH 7.5, 5 mM MgCl_2_, 1 mM ATP, 10% glycerol, 0.2 μg/μl BSA, 1 mM DTT, and 1 µl Rnasein (Promega), in presence of increasing concentration (25-400 nM) of FMRP-WT, FMRP-I1034N, N-Fold-WT, N-Fold-I304N or C-IDR, as indicated. The reactions were stopped with addition of 1 μl 1% SDS and 1 μl 10 mg/ml Proteinase K (Invitrogen™), and incubating at 37°C for further 5 m. Finally, 2 µl loading buffer composed of 50% glycerol, 20 mM Tris-HCl, pH 7.4, 0.5 mM EDTA, 0.05% Orange G, was added to each tube, and the products were resolved by running in 10% TAE gel, at 100V for 60 minutes. The gels were dried, exposed to phosphorimaging screen, and imaged as discussed earlier.

### DNA:RNA immunoprecipitation (DRIP)

12-18×10^6^ cells were treated with DMSO and APH or left untreated. 24 hr post treatment, cells were washed twice with 1X PBS and centrifuged at 250 g for 5 m at 4°C. Pellets were quick-frozen and stored in −70°C until use. Cell pellets were thawed on ice. Genomic DNA and DNA:RNA hybrid were isolated by adding 5 ml 1X lysis buffer (50 mM Tris-HCl pH 8.0/1M NaCl/10 mM EDTA/0.5% SDS/0.2 mg/ml Proteinase K) prewarmed at 37°C to the thawed cells. The mixture was carefully inverted a few times and incubated overnight at 37°C. Nucleic acids were purified using phenol: chloroform: isoamyl alcohol and ethanol precipitation. Upon precipitation DNA was spooled or centrifuged at 3000 rpm for 10 m. DNA was washed with 70 % alcohol, air-dried and resuspended in elution buffer. DNA was sonicated in Covaris M220 ultrasonicator using AFA fiber pre-slit snap-cap 130 μl microtubes and in-built protocol for fragmentation of DNA to 300-500 bp. 16 μg of sonicated DNA was subjected to RNase H (NEB, 1.5 U per μg of DNA) treatment or RNase H buffer only for 4 hr at 37°C. 10% of the reaction was aliquoted and saved as ‘Input’, while the rest was used for immunoprecipitation. Immunoprecipitation was carried out in 1X DRIP binding buffer (0.02% Tween 20 in 1X PBS) with 11 μg of S9.6 antibody (Kerafast) per reaction overnight at 4°C with constant rotation. Sixty microliter of Dynabead protein G (Invitrogen) was washed twice in 1X DRIP binding and added to the immuno-complex, incubated for 2 hr at 4°C with constant rotation. Beads were washed with 750 μl of 1X DRIP binding buffer twice, followed by elution in 300 μl of DRIP elution buffer (50 mM Tris-HCl pH 8.0/10 mM EDTA pH 8.0/0.5% SDS in DEPC water) and 7 μl of 20 mg/ml Proteinase K. The immuno-complex was eluted twice at 55°C for 45 m under constant rotation. The ‘Input’ samples and the eluted immune complex was then purified with phenol: chloroform: isoamyl alcohol and ethanol precipitation.

### Chromatin immunoprecipitation (ChIP)

#### Cell collection

FMR1KO-B3-EGFP, FMR1KO-B3-EGFP-FMRP-H9 and FMR1KO-B3-EGFP-FMRPI304N-F10 cells were grown to 75% confluency. Cells were treated with DMSO/APH or left untreated for 24 hrs. Cells were fixed and harvested using the truChIP chromatin shearing kit (Covaris) and IP was conducted according to Richard Myers lab ChIP-seq protocol. Briefly, cells were first washed with room temperature 1X PBS and then 5 ml fixing buffer A was added. 500 μl of 11.1% formaldehyde(methanol-free) was added to the cells and incubated for 10 m at room temperature. To stop the reaction, 300 μl of quenching buffer was added to the mix for 5 m. Cells were scrapped and collected (one confluent 150 mm plate per reaction) frozen in liquid nitrogen and stored at −70°C until further use. Cells were thawed in ice with 1 ml of 1X lysis buffer B and Halt protease inhibitor cocktail (Thermo scientific)] and incubated in a rotator for 10 m. Nuclei were prepared by centrifuging the lysate at 1700 g for 5 m. The nuclear pellet was washed with 1X wash buffer C and then once with shearing buffer D3 with Halt protease inhibitor cocktail (Thermo scientific)]. Nuclear pellet was then resuspended in 900 μl shearing buffer D3 and sonicated in Covaris M220 ultrasonicator using the in-built protocol ‘ChIP_10%df_10min’ (75W peak power, cycle per burst : 200, duty factor: 10, time: 600 s. The sonicated mixture was centrifuged at 13,500 rpm for 15 m at 4°C. The supernatant was diluted to adjust the salt concentration to 150 mM NaCl, 10 mM Tris-HCl pH 7.5, 1% NP-40, 0.5% sodium deoxycholate. Approximately 5-11% of this mixture was set aside as ‘input’. The rest was used for immunoprecipitation. Both the aliquots were snap frozen in liquid nitrogen and stored at −70°C.

#### Immunoprecipitation

300 μl M280 Dyna beads sheep anti-rabbit IgG (Life technologies) was added to 1 ml freshly prepared PBS with 5 mg/ml BSA (PBS/BSA) and Halt protease inhibitor cocktail (1X). The magnetic beads were washed 3 times in PBS/BSA. 12 μg of monoclonal anti-DHX9 antibody (Bethyl Laboratories) or 15 mg of FMRP (Biolegend) antibody was mixed with the beads in 1 ml PBS/BSA and incubated overnight at 4°C with agitation. Next day, the antibody solution was removed and the beads were washed 3 times with PBS/BSA. The sonicated nuclear fraction for each sample was thawed and added to one-third of the beads prepared and mixed well. The samples were then incubated overnight at 4°C with rotation. The following day, beads were washed 5 times with cold LiCl wash buffer (100 mM Tris pH 7.5, 500 mM LiCl, 1% NP-40, 1% sodium deoxycholate). This was followed by a single wash in TE buffer (10 mM Tris-HCl pH 7.5, 0.1 mM EDTA). 200 μl of IP elution buffer (1% SDS, 0.1 M NaHCO_3_) was added to the beads, mixed and incubated at 65°C for 2 h with vortex every 30 m. The ‘input’ was thawed and together with the supernatant from the immunoprecipated sample was reverse-crosslinked at 65°C overnight. Post reverse cross-linking, samples were treated with 60 μg of proteinase K for 1 hr at 55 °C DNA was purified using Qiagen PCR purification kit. The purified DNA was then used for qPCR.

### Quantitative Real-Time PCR (qPCR) and PCR

After IP and DNA isolation, the ChIP’ed and DRIP’ed DNA and the input DNA was diluted in water. The PCR reaction was carried out in 10 μl volume with 5 μl of 2X iTaq Universal Sybergreen Supermix (BioRad), 500 nM of forward and reverse primers (for promoter, intron-5 and pause sites from Cristini et al, 2018), 3 μl of DNA and water. PCR conditions were obtained from Sanz et al, 2019. Briefly, CFX Opus 384 real time PCR system (BioRad) was used for qPCR with the following thermal cycling protocol: 1 cycle of 95°C for 30 s; 39 cycles of 95°C for 10 s and 60°C for 30 s; followed by melt curve analysis from 65°C to 95°C by an increment of 0.5°C for 5 s. qPCR data was analyzed and “% Input” was calculated as described (Sanz and Chédin, 2019). Sonicated genomic DNA was used to amplify the β-actin locus using forward and reverse primers for promoter, intron-5 and pause sites from Cristini et al. (Cristini et al., 2018). A reaction volume of 25 μl containing 1XPrimeStar Max DNA Polymerase (Takara), 300 nM of forward and reverse primers and 100 ng of DNA. BioRad MyCycler with the following thermal cycling protocol: 1 cycle of 98°C for 30 s; 30 cycles of 98°C for 10 s and 60°C for 30 s was used. Products were run on a 2% agarose gel in Tris-Borate EDTA buffer with 0.3 mg/ml ethidium bromide concentration.

## SUPPLEMENENTARY INFORMATION

### FIGURE SUPPLEMENTS

**Figure S1.**
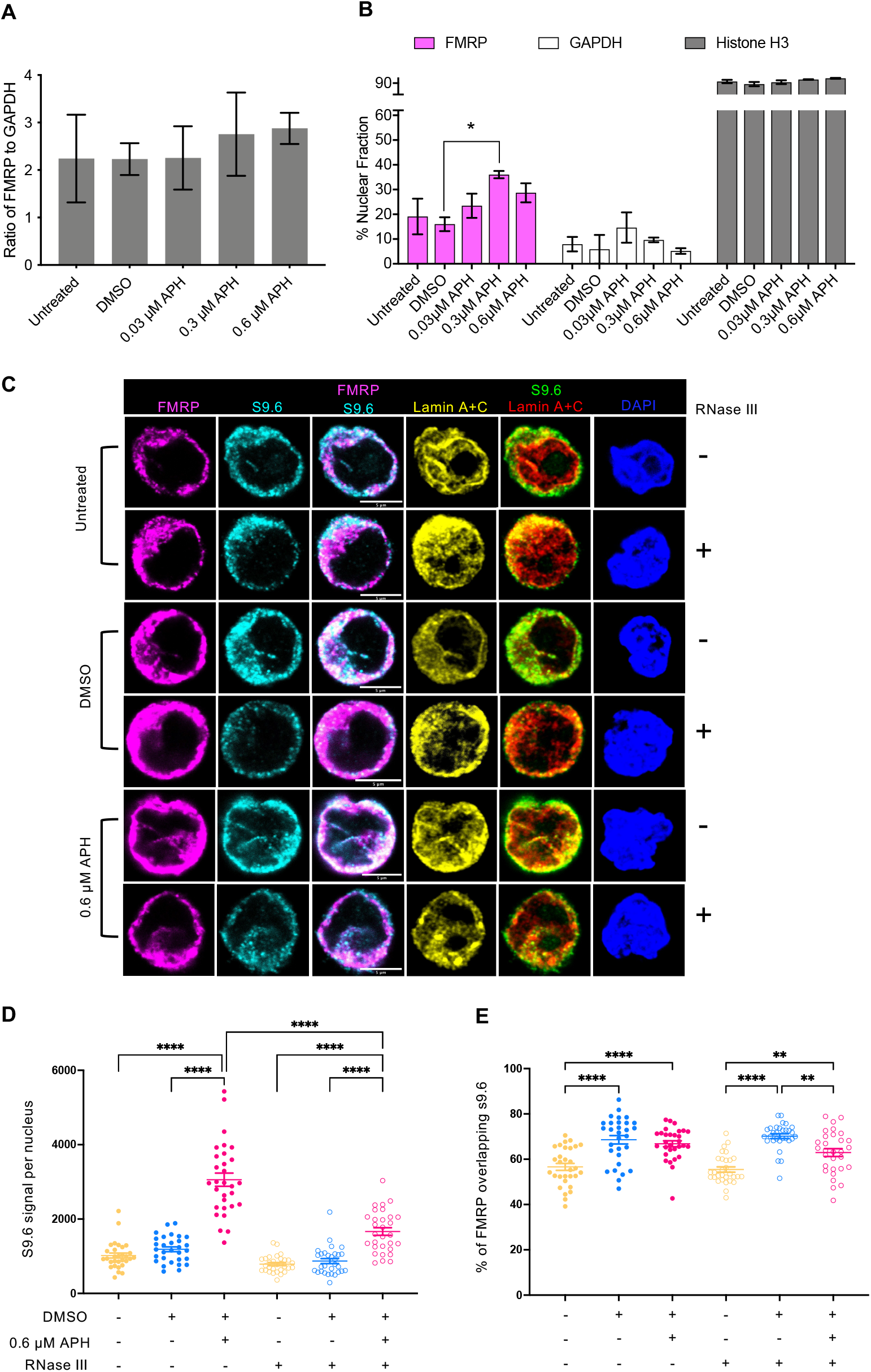
**(A)** Total FMRP level expressed as ratio of FMRP over GAPDH in the whole cell extracts (n=2) from Figure 1A remained nearly constant in all conditions. **(B)** Quantification of FMRP, GAPDH and Histone H3 intensity shows increased percentage of FMRP in the nuclear fraction under APH stress. Percentage of nuclear fraction of proteins expressed as the percentage of the band intensity for “N” over that of the sum of “N” and “C” for each condition. Error bars indicate standard error of mean in two independent experiments. One-way ANOVA followed by Tukey’s multiple comparison test. *, p = 0.033. **(C)** Co-localization of FMRP and RNA:DNA hybrids. Immunofluorescence images of untreated, DMSO and APH treated GM6990 cells co-stained for RNA:DNA hybrids (cyan), FMRP (magenta), lamin A/C (yellow) and DAPI (blue). Cells were treated with RNase III enzyme to show specificity for staining R-loops by S9.6 antibody. Immuno-staining is shown in a single Z-plane. Scale bar, 5 µm. **(D)** Quantification of S9.6 signal per nucleus in cells treated with or without RNase III. Error bars indicate SEM, N∼30 cells. One-way ANOVA followed by Tukey’s multiple comparison test, **p<0.01, ****p < 0.0001. **(E)** Quantification of colocalization of FMRP with S9.6 signal using Coloc2 plugin in Fiji. Error bars indicate SEM, N∼30 cells per sample. One-way ANOVA followed by Tukey’s multiple comparison test, *p < 0.05, **p<0.01, ****p < 0.0001

**Figure S2.**
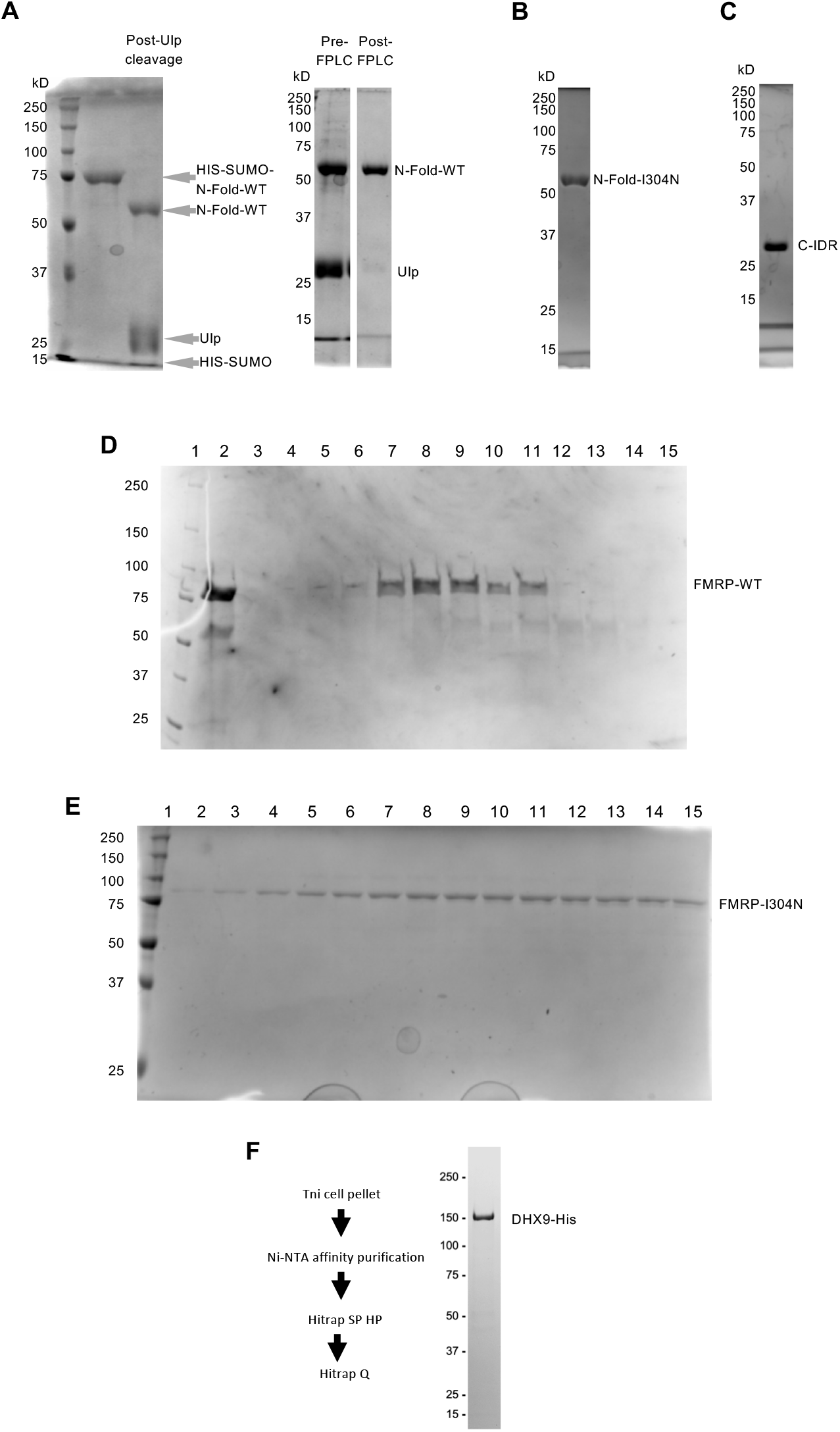
Purification of FRMP fragments, their interactions with various nucleic acid structures and purification of DHX9. **(A-C)** Purification of FMRP protein domains for EMSA. The fusion proteins containing HIS-SUMO-tagged FMRP fragments were subject to Ulp cleavage to remove the tag, followed by FPLC to remove the cleaved HIS-SUMO as well as Ulp itself, as shown for N-Fold-WT (A). The same procedures were applied to the purification of N-Fold-I304N (B) and C-IDR (C). (**D&E**) Representative SDS-PAGE gel images of FPLC gel filtration column purification of FMRP-WT (D) and FMRP-I304N (E). Lanes in D are: 1, ladder; 2, input/load; 3-15, fractions from the FPLC Gel Filtration purification; 7-9, fractions contain pure FMRP and were collected and combined; 3-6 and 10-15 fractions were discarded due to low FMRP concentration and/or the presence of degradation products. Lanes in E are: 1, ladder; 2-15 fractions from the FPLC Gel Filtration purification. (**F**) Purification of DHX9-His.

**Figure S3.**
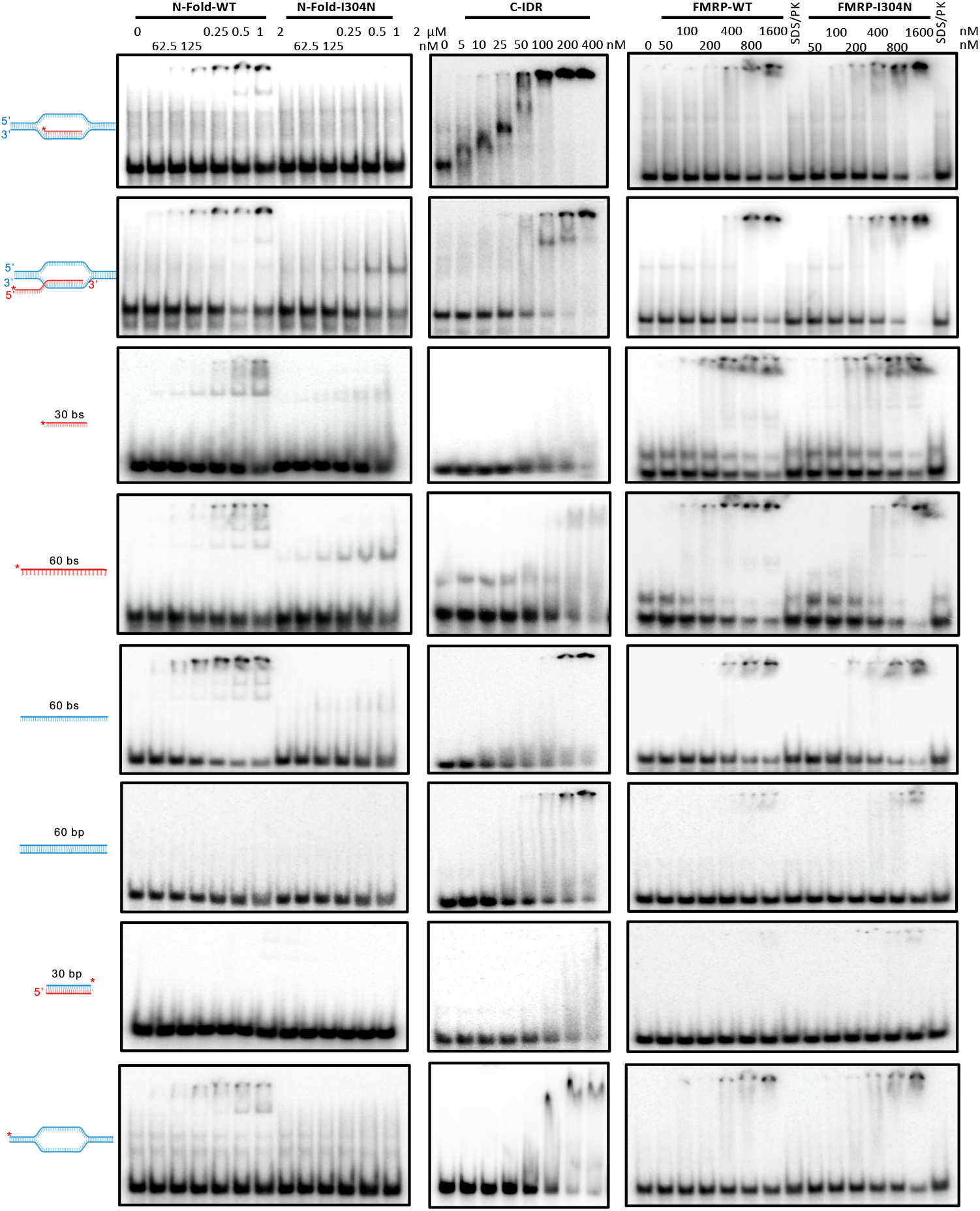
Representative EMSA for all proteins and nucleic acids to accompany results in Fig. 2.

**Figure S4.**
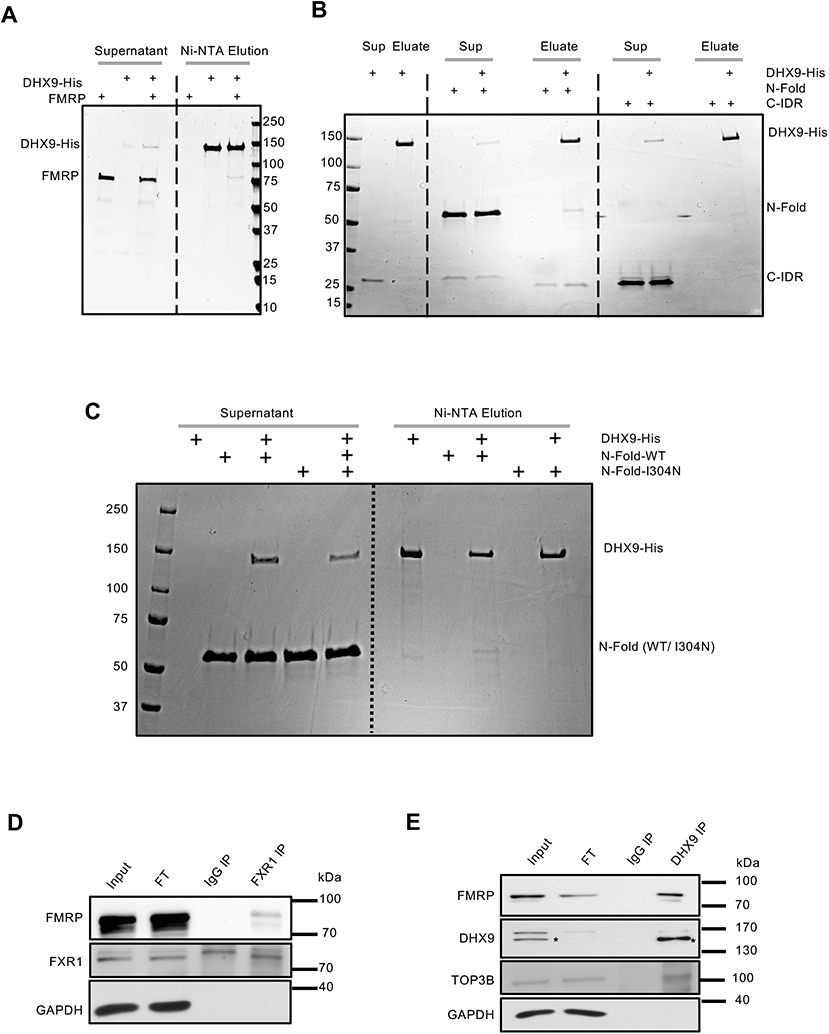
FMRP interacts with DHX9 *in vitro* and *in vivo.* **(A-C)** *In vitro* protein binding assays for DHX9-His and full length FMRP **(A)**, FMRP domains **(B)** and N-Fold-WT or N-Fold-I304N **(C). (D)** Co-immunoprecipitation of FMRP by immunoprecipitating with anti-FXR1 monoclonal antibody and immunoblotted for FMRP and FXR1. GAPDH served as negative control. **(E)** Co-immunoprecipitation of FMRP by immunoprecipitating with anti-DHX9 monoclonal antibody and immunoblotted for FMRP, DHX9 and TOP IIIβ. The black asterisks indicate the lower band of a doublet signal in the “IP-DHX9” lane is the DHX9 protein, which is accumulated in the immunoprecipitated complex and absent in the IgG-precipitated control complex (“IP-IgG” lanes).

**Figure S5.**
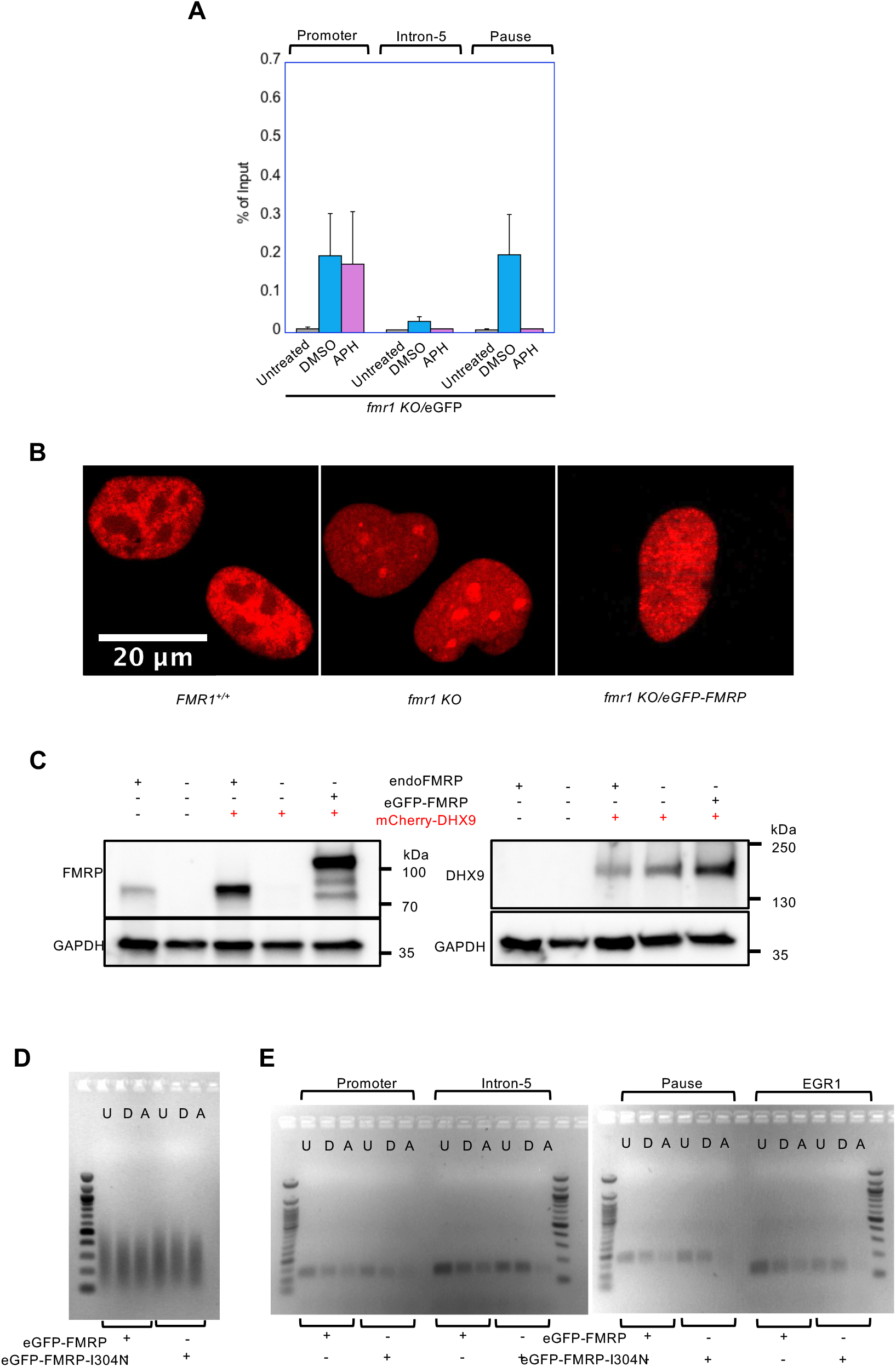
Functional interaction between FMRP and DHX9. **(A)** DHX9 ChIP-qPCR across the β-actin locus from *fmr1 KO* cells expressing eGFP only. Two independent experiments were done and for each replicate and percentage of input was calculated. The average of the two experiments calculating percentage of input is shown. **(B)** Altered nuclear localization of mCherry-DHX9 in cells lacking FMRP. Live cell images of cells overexpressing DHX9 with mCherry tag in the indicated background. **(C)** Western blots of the cells shown in (B) probed for FMRP (left) or mCherry (right). The first two lanes in both western blots are control cell lines. “endoFMRP”, endogenous FMRP. **(D)** Agarose gel image of sonicated genomic DNA isolated from eGFP-FMRP and eGFP-FMRP-I304N expressing cells in untreated-U, DMSO-D and APH-A treated conditions. These samples were used for DRIP in Figure 5(C). **(E)** PCR amplification of β-actin locus from sequences in Figure 5 and using DNA template from (D).

**Table S1.**
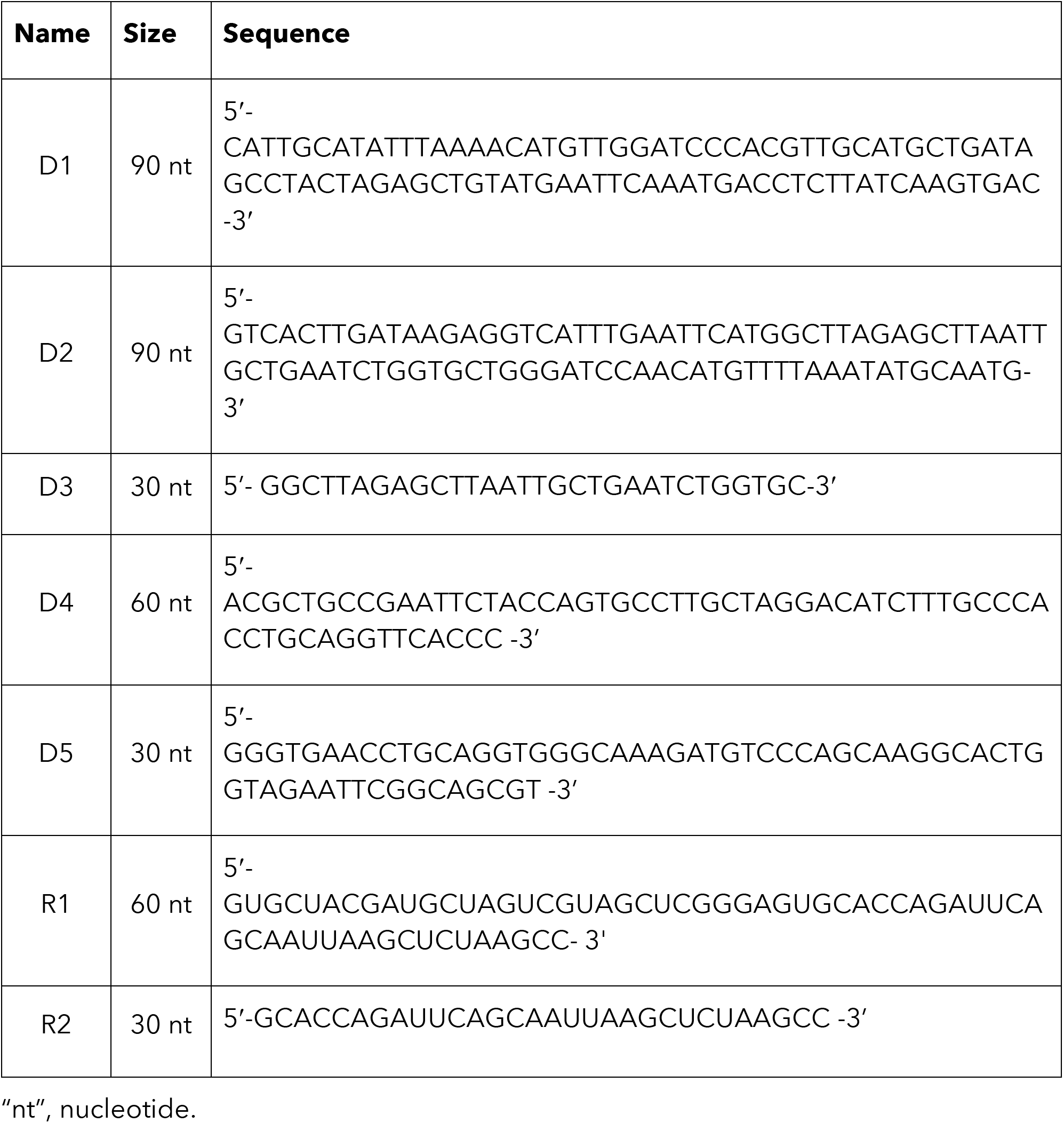
List of all oligonucleotides for making substrates for EMSA experiments.

**Table S2.** Scheme for generating substrates for EMSA experiments.

|  |  |
| --- | --- |
| R-loop with 5'-RNA ovh | D1+D2+R1 |
| R-loop with no-RNA ovh | D1+D2+R2 |
| RNA-DNA hybrid with 5'-RNA ovh | D3+R1 |
| RNA-DNA hybrid with no RNA ovh | D3+R2 |
| dsDNA | D4+D5 |
| Bubble DNA | D1+D2 |
| ssDNA | D1 |
| RNA (60 nt) | R1 |
| RNA (30 nt) | R2 |

"ovh", overhang.

